# Small Molecule Inducers of Neuroprotective miR-132 Identified by HTS-HTS in Human iPSC-derived Neurons

**DOI:** 10.1101/2022.11.01.514550

**Authors:** Lien D. Nguyen, Zhiyun Wei, M. Catarina Silva, Sergio Barberán-Soler, Rosalia Rabinovsky, Christina R. Muratore, Jonathan M. S. Stricker, Colin Hortman, Tracy L. Young-Pearse, Stephen J. Haggarty, Anna M. Krichevsky

**Author notes:** Lien D. Nguyen and Zhiyun Wei contributed equally to this work.

## Abstract

MicroRNAs (miRNAs) are short RNAs that regulate fundamental biological processes. miR-132, a key miRNA with established functions in Tau homeostasis and neuroprotection, is consistently downregulated in Alzheimer’s disease (AD) and other tauopathies. miR-132 overexpression rescues neurodegenerative phenotypes in several AD models. To complement research on miRNA-mimicking oligonucleotides targeting the central nervous system, we developed a high-throughput-screen coupled high-throughput-sequencing (HTS-HTS) in human induced pluripotent stem cell (iPSC)-derived neurons to identify small molecule inducers of miR-132. We discovered that cardiac glycosides, which are canonical sodium-potassium ATPase inhibitors, selectively upregulated miR-132 in the sub-μM range. Coordinately, cardiac glycoside treatment downregulated total and phosphorylated Tau in rodent and human neurons and protected against toxicity by glutamate, N-methyl-D-aspartate, rotenone, and Aβ oligomers. In conclusion, we identified small-molecule drugs that upregulated the neuroprotective miR-132 and ameliorated neurodegenerative phenotypes. Our dataset also represents a comprehensive resource for discovering small molecules that modulate specific miRNAs for therapeutic purposes.

## INTRODUCTION

Despite the enormous burden of Alzheimer’s disease and related dementias (ADRDs) on patients, caregivers, and society, there is still a lack of effective, disease-modifying treatments. Traditional drug discovery has focused on disease-relevant proteins and peptides such as Aβ, Tau, and β-amyloid cleaving enzyme ^1^. However, RNAs have recently emerged as promising targets for broad disease categories, with several approved RNA therapeutics in the last five years ^2^. Particularly, >70% of the human genome is transcribed into noncoding RNAs (ncRNAs) that play essential yet largely understudied roles in biological processes ^3, 4^. MicroRNAs (miRNAs) are short, single-stranded ncRNAs of 18–25 nucleotides that facilitate the degradation and inhibit the translation of mRNA targets ^5^. Specific miRNAs have been shown to be dysregulated in various diseases ^6^, making them valuable targets for both diagnostic and therapeutic purposes.

Here, we focus on miR-132, one of the most consistently downregulated miRNAs in the cortex and hippocampus of ADRD patients ^7–11^. miR-132 deficiency promotes Aβ plaque deposits ^12, 13^, and Tau accumulation, phosphorylation, and aggregation ^13–16^. We recently showed that miR-132 mimics protected mouse and human primary neurons against Aβ oligomer and glutamate toxicity^16^. miR-132 viral overexpression reduced Tau toxicity and neuronal loss in presymptomatic PS19 mice overexpressing an autosomal dominant mutation (P301S) in the human microtubule-associated protein tau (*MAPT*) transgene, likely through directly targeting Tau modifiers, including glycogen synthase kinase 3 β (*GSK3β*), E1A binding protein P300 (*EP300*), RNA binding Fox-1 homolog 1 (*RBFOX1*), and calpain-2 (*CAPN2*) ^16^. Furthermore, miR-132 level was decreased in the hippocampus of several AD mouse models, and viral overexpression of miR-132 rescued adult hippocampal neurogenesis and memory deficits in these models ^11, 12, 16, 17^. These findings collectively support upregulating miR-132 in the central nervous system (CNS) as a promising approach for preventing or treating AD and tauopathies.

Two common approaches to upregulating miRNAs, oligonucleotide mimics and gene delivery, have serious limitations. miRNA mimics, which are synthetic oligonucleotides imitating mature miRNA sequences and structures, often have poor intracellular delivery and on-target activity, must be heavily modified to avoid rapid degradation, and can induce immunotoxicity ^18–20^. Similarly, delivering genes coding for miRNAs through viral or non-viral vectors is generally inefficient and can induce immunotoxicity or off-target integration ^19^. The CNS presents additional challenges for drug delivery and efficacy due to the blood-brain barrier that blocks the entrance of most compounds. We proposed small molecules as an alternative approach for upregulating miRNAs ^21^. Compared to miRNA mimics and gene therapy, small molecules usually have better brain and cell penetrance. Small molecules already approved for treating human diseases have well-established safety profiles and pharmacokinetics. Repurposing or improving these compounds would accelerate the development of miRNA therapeutics to enter clinical trials. However, only a few small molecules affecting miRNA levels have been described ^22^, and no systematic effort has been made to identify such modulators of miRNA expression and activity. Therefore, we designed a pipeline for discovering small molecules that upregulate miR-132 in human induced pluripotent stem cell (iPSC)-derived excitatory neurons. To our knowledge, no miRNome-wide high-throughput screen of small molecule modulators of miRNA, and particularly miRNA inducers, has been developed to date.

We performed high-throughput-screen coupled high-throughput-sequencing (HTS-HTS) of ∼1900 bioactive compounds in iPSC-derived human neurons and validated that several members of the cardiac glycoside family, which are sodium-potassium (Na+/K+) ATPase pump inhibitors, upregulated miR-132 in the sub-μM range. Treating rodent and human neurons with sub-μM cardiac glycosides protected neurons against various toxic insults and downregulated Tau and other miR-132 targets. Overall, we identified small-molecule compounds that upregulated the neuroprotective miR-132 in neurons and provided a pipeline for discovering small-molecule compounds that regulate miRNAs for therapeutic purposes.

## RESULTS

### Optimization of the high-throughput screen on human iPSC-derived neurons

We used human neurogenin 2 (NGN2)-driven iPSC-derived neurons (NGN2-iNs) as a physiologically relevant cell-based screening platform to discover miR-132 inducers. iPSC lines generated from donors were utilized for direct differentiation through NGN2 overexpression into excitatory neurons based on established protocols (Figure 1A) ^23^. These cells closely mimic the transcriptome and function of human neurons *ex vivo* and can be scaled and reproducibly employed in multiple assays ^23^. Among 36 NGN2-iN lines obtained from the Religious Orders Study/Memory and Aging Project (ROS-MAP) cohort, 25 lines from donors without cognitive impairment were considered (S1A). The transcriptomes of these lines were previously profiled ^23^. The BR43 line was selected for the screen based on its median expression of major miR-132 targets, including *GSK3β, EP300, RBFOX1, CAPN2, FOXO3, TMEM106B, and MAPT* (S1B). Importantly, BR43 NGN2-iNs had the lowest variation of baseline miR-132 expression among the replicate cultures and exhibited miR-132 upregulation by the known inducers BDNF and forskolin, thus providing a reliable platform for the high-throughput screen (S1C).

**Figure 1.**
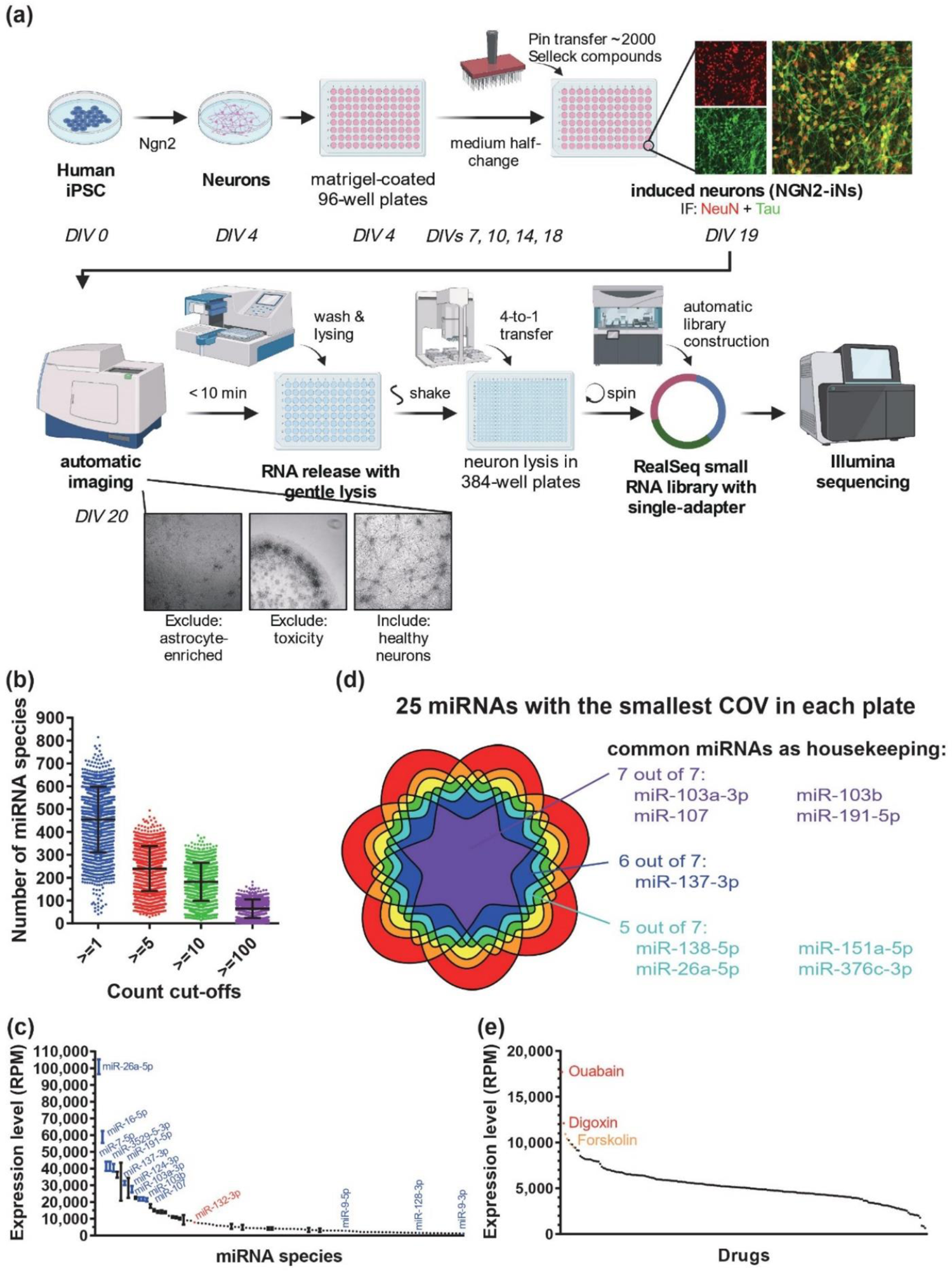
Experimental workflow and overview of screen results. **a**, NGN2-iN generation, drug treatment, and miRNA-seq workflow (N=1 per drug). **b**, Average number of miRNA species detected per sample by miRNA-seq at various count cut-offs. **c**, Expression levels of the 100 most abundant miRNAs in vehicle-treated samples. miR-26a-5p was the most abundant miRNA detected, and miR-132-3p was the 27^th^. **d**, Shared miRNAs with the lowest coefficient of variation among the 7 plates tested. **e**, Waterfall plot for miR-132 expression in plate 2. Samples treated with ouabain, digoxin, and the positive control forskolin showed the highest level of miR-132.

Several steps of NGN2-iN culture and RNA collection were optimized for the high-throughput screen (HTS) to maximize neuronal health, lysing efficiency, and RNA yield (S1D, E). The protocol was tested for its compatibility with small RNA-seq using the RealSeq ultra-low input system, long RNA RT-qPCR using the PrimeScript system, and small RNA RT-qPCR using the miRCURY system (S1D, E), further supporting its application in diverse quantitative RNA-based assays.

### Small molecules screen of miR-132 inducers

Day 4 NGN2-iNs were plated onto 25 Matrigel-coated 96-well plates and differentiated into neurons, as verified by NeuN and Tau expression (Figure 1A). On day 19, the Selleckchem library (N=1,902 compounds), a diverse library of bioactive molecules, was pin-transferred into plates to achieve 10 μM final concentration. DMSO (0.1% final concentration) and forskolin (10 μM) were used as the negative and positive controls, respectively. NGN2-iNs were imaged to monitor neuronal health 24h later, followed by direct lysis to release RNA (Figure 1A). Among all wells with test compounds, 324 (17.0%) were excluded because of cell death, neurite degeneration, loss of cells during washes, or enrichment of astrocytes. RNA lysates of the remaining wells, including positive and negative controls, were used for RealSeq small RNA library preparation designed for ultra-low input without RNA purification ^24^. RealSeq libraries from each set of four 96-well culture plates were indexed with 384 multiplex barcodes and pooled for deep sequencing (Figure 1A). After miRNA annotation, wells with less than 1,000 total annotated read counts were excluded from further analysis (N=169, 10.7%). On average, 55,529 miRNA reads were counted per sample, and 455, 240, 182, and 64 miRNA species per sample were detected with minimal read counts of 1, 5, 10, and 100, respectively (Figure 1B). Numerous neuron-enriched miRNAs, such as miR-124, miR-26a, miR-128, miR-9, and miR-191, were abundant in DMSO-treated control NGN2-iNs (Figure 1C). As expected, miR-132 was consistently detected and ranked among the 30 most abundant miRNAs (Figure 1C). We further determined the top housekeeping neuronal miRNAs by calculating the coefficient of variation (COV) for each miRNA within each batch of RNA-seq and identified the miRNAs with the smallest COVs, including miR-103a/b, miR-107, and miR-191 (Figure 1D). As library preparation and sequencing for different 384-well plates were carried out on different days, to minimize batch effects, compounds within each 384-well plate were ranked for miR-132 expression. Figure 1E showed the miR-132 waterfall plot for 221 compounds in a 384-well plate.

### Selection of compounds for further validation

To select compounds for further validation, we used a matrix with miR-132 plate rank as the primary criterion and adjusted with secondary criteria, including the US Food and Drug Administration (FDA) approval, BBB penetrance, clinical trials, published data on neuroprotective effects, and effects on other miRNAs (Table S1). We treated DIV14 primary rat cortical neurons and DIV21 human NGN2-iNs with 10 μM of 44 reordered compounds, and monitored miR-132 expression by RT-qPCR. 12 and 10 compounds significantly upregulated miR-132 in rat neurons after 24h and 72h, respectively, and 4 compounds significantly upregulated miR-132 in NGN2-iNs after 24h (Figure 2A, Table S2). Notably, the cardiac glycosides, ouabain and digoxin, upregulated miR-132 in all conditions. Several chemotherapeutics, including rigosertib, pelitinib, letrozole, XL888, and etoposide, appeared to mildly upregulate miR-132 but also caused toxicity after 72h.

**Figure 2.**
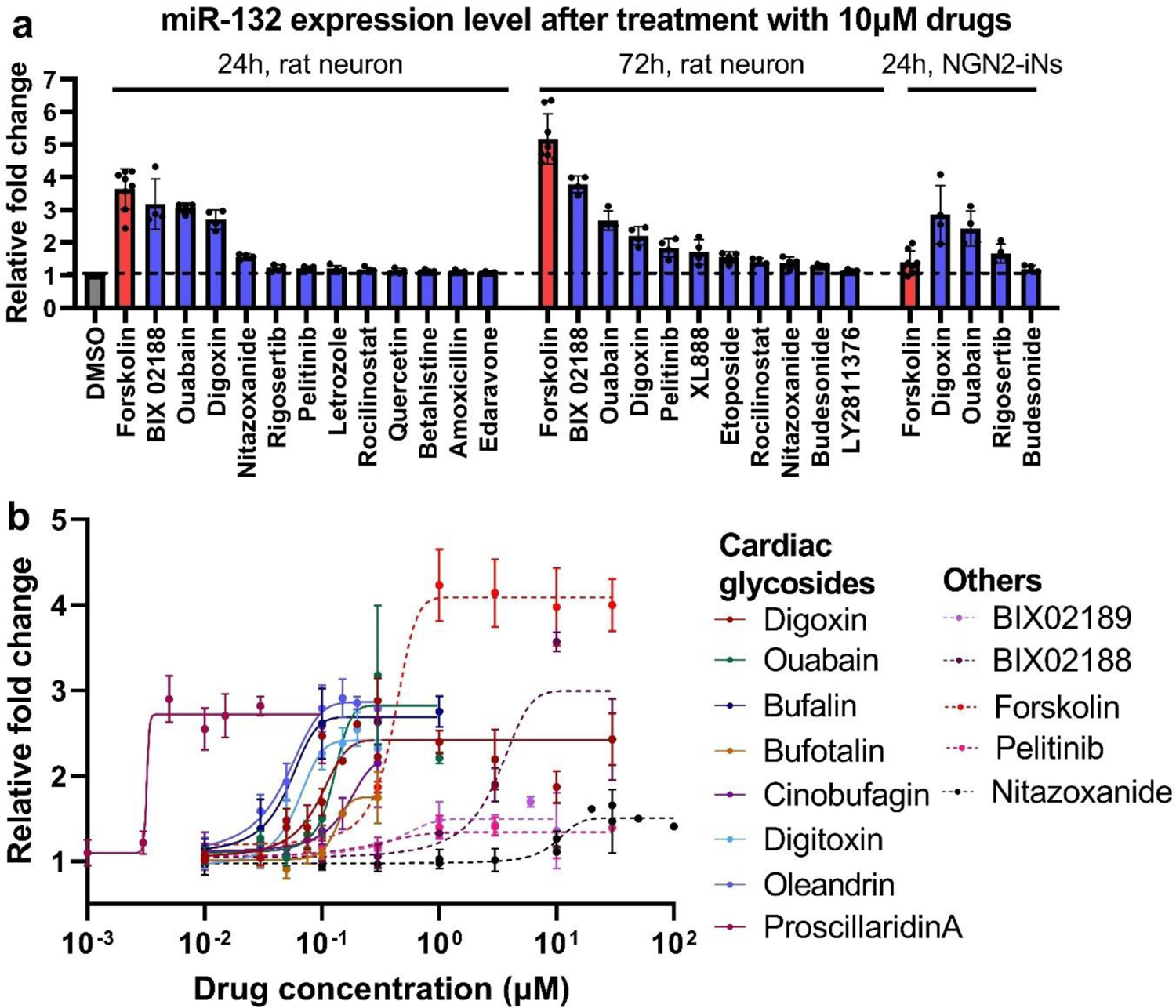
Validation of top candidates from HTS-HTS and dose curve experiment. **a**, Drugs that showed significant upregulation of miR-132 in primary rat cortical neurons after 24 and 72h treatment and human NGN2-iNs after 24h treatment (RT-qPCR analysis, N=4, unpaired two-tailed Student’s t-test, p<0.05 compared to DMSO). **b**, Dose curve experiments were performed in DIV14 rat neurons after 24h treatment. Solid lines were used for cardiac glycosides, and dotted lines were used for other drugs. EC_50_ and max fold change were calculated using sigmoidal fit, 4 parameters. (N=4-6, error bars represent SD).

### Dose-response of hit compounds

To investigate dose response, we selected forskolin as the positive control, digoxin, ouabain, BIX02188, nitazoxanide as the hits, and pelitinib as a representative chemotherapeutic. We also included 6 additional cardiac glycosides (digitoxin, oleandrin, bufalin, bufotalin, cinobufagin, and proscillaridin A) and BIX02189, an analog of BIX02188. These compounds represent diverse chemical groups and mechanisms of action (Figure 2B and Table S3). DIV14 primary rat cortical neurons were treated with drugs at doses ranging from 1 nM to 100 μM for 24h. Remarkably, all 8 cardiac glycosides upregulated miR-132 2.5-3-fold in the sub-μM range, with proscillaridin A having the lowest EC_50_ of 3.2 nM (Table S3). Other compounds also dose-dependently upregulated miR-132 but with higher EC_50_. For all compounds tested, miR-212, which is a miRNA co-transcribed and co-functional with miR-132 ^25^, was similarly upregulated at almost identical EC_50_, suggesting that the mechanism was largely transcriptional (S2A and Table S3). The cardiac glycosides proscillaridin A, oleandrin, digoxin, ouabain, and bufalin also upregulated miR-132 and miR-212 in a dose-dependent manner in human NGN2-iNs in the sub-μM range (S2B, C and Table S3). However, BIX02188, which robustly upregulated miR-132 in primary rat neurons, had no effect on miR-132 in NGN2-iNs (S2C and Table S3), suggesting potential differences between the two cell models.

### Upregulation of miR-132/212 was specific and transcriptional

To investigate the specificity of miR-132 upregulation, we measured the expression level of 10 other abundant neuronal miRNAs in rat primary cortical neurons after 24h of treatment with oleandrin and BIX02188. When normalized to the geometric mean of all 12 miRNAs ^26^, only miR-132 and miR-212 were upregulated (Figure 3A). The precursors pre-miR-132 and pre-miR-212 (Figure 3B) were also upregulated by forskolin, BIX02118, and the cardiac glycosides, suggesting that these compounds activated the transcription of the miR-132/212 locus. Correspondingly, the upregulation of miR-132 by forskolin and oleandrin was completely blocked by pretreatment with the transcription inhibitor actinomycin D (Figure 3C, S3). As miR-132/212 locus is regulated by the transcription factor CREB ^27^, cells were also pretreated with the maximum tolerated doses of a CREB inhibitor (1 μM CREB-I). Co-treatment with the CREB inhibitor partially attenuated miR-132 upregulation by forskolin and oleandrin (Figure 3C). As cardiac glycosides are conventional inhibitors of Na^+^/K^+^ pumps, we also knocked down ATP1A1 and ATP1A3, the dominant isoforms in neurons, with siRNAs. Knocking down either ATP1A1 or ATP1A3 also increased the expression of products of the miR-132/212 locus (Figure 3D), suggesting that cardiac glycosides upregulated miR-132 by inhibiting their conventional targets.

**Figure 3.**
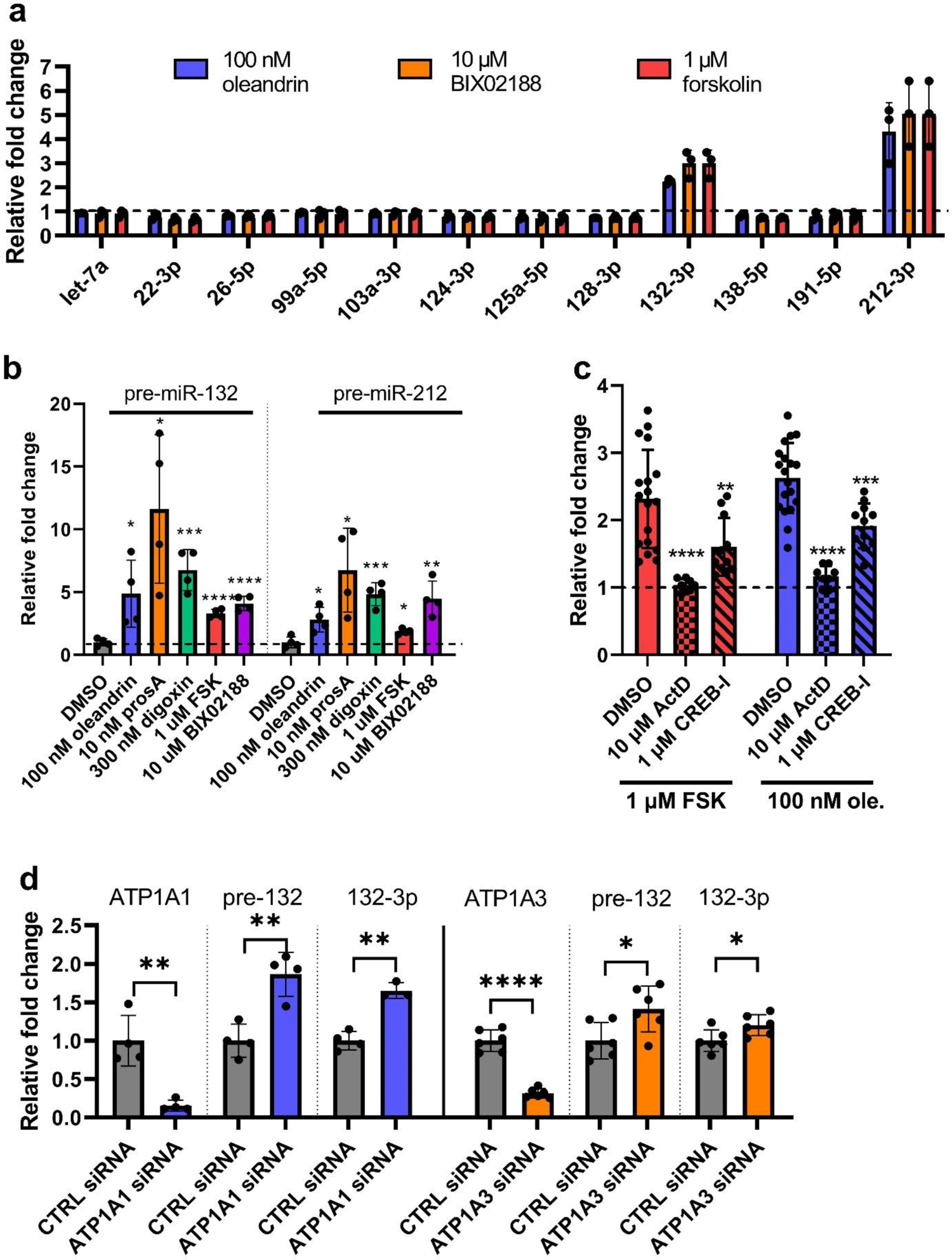
Cardiac glycosides transcriptionally upregulated miR-132/212 by inhibiting the Na^+^/K^+^ ATPases. **a**, Forskolin, oleandrin, and BIX02188 specifically upregulated miR-132/212 without affecting other abundant miRNAs. Expression was normalized to the geometric mean of all 12 miRNAs tested. **b**, Cardiac glycosides, forskolin, and BIX02188 also upregulated the precursors of miR-132/212 24h after treatment (unpaired two-tailed Student’s t-test compared to DMSO control, N=4). **c**, Upregulation of miR-132 by forskolin or oleandrin was completely blocked by the transcription inhibitor actinomycin D and partially blocked by CREB inhibitor (unpaired two-tailed Student’s t-test compared to DMSO control, N=8-19). **d**, Knocking down ATP1A1 or ATP1A3, the predominant isoforms in neurons, also upregulated pre- and mature miR-132 (unpaired two-tailed Student’s t-test compared to DMSO control, N=4-6).

### Kinetics of miR-132 upregulation and effects on known targets

To investigate the kinetics of miR-132 upregulation by cardiac glycosides, we treated primary rat cortical neurons with 100 nM oleandrin and measured the expression of the precursor and the mature forms of miR-132 and miR-212 overtime (Figure 4A, B). Both pre-miR-132 and pre-miR-212 were rapidly upregulated following treatment, peaked at 8h, and rapidly declined back to baseline after 72h (Figure 4A). As oleandrin was shown to upregulate *BDNF* ^28, 29^, a known transcriptional regulator of the miR-132/212 locus ^27^, we also measured *BDNF* expression level. Oleandrin upregulated *BDNF* as expected but at a slower kinetics than pre-miR-132/212, suggesting that *BDNF* was not mediating the effects of oleandrin on miR-132/212 expression (Figire 4A). Compared to their precursors, mature miR-132 and −212 were upregulated at slower kinetics, peaked at 24h, then slowly declined but were still ∼2-fold above baseline at 72h (Figure 4B).

**Figure 4.**
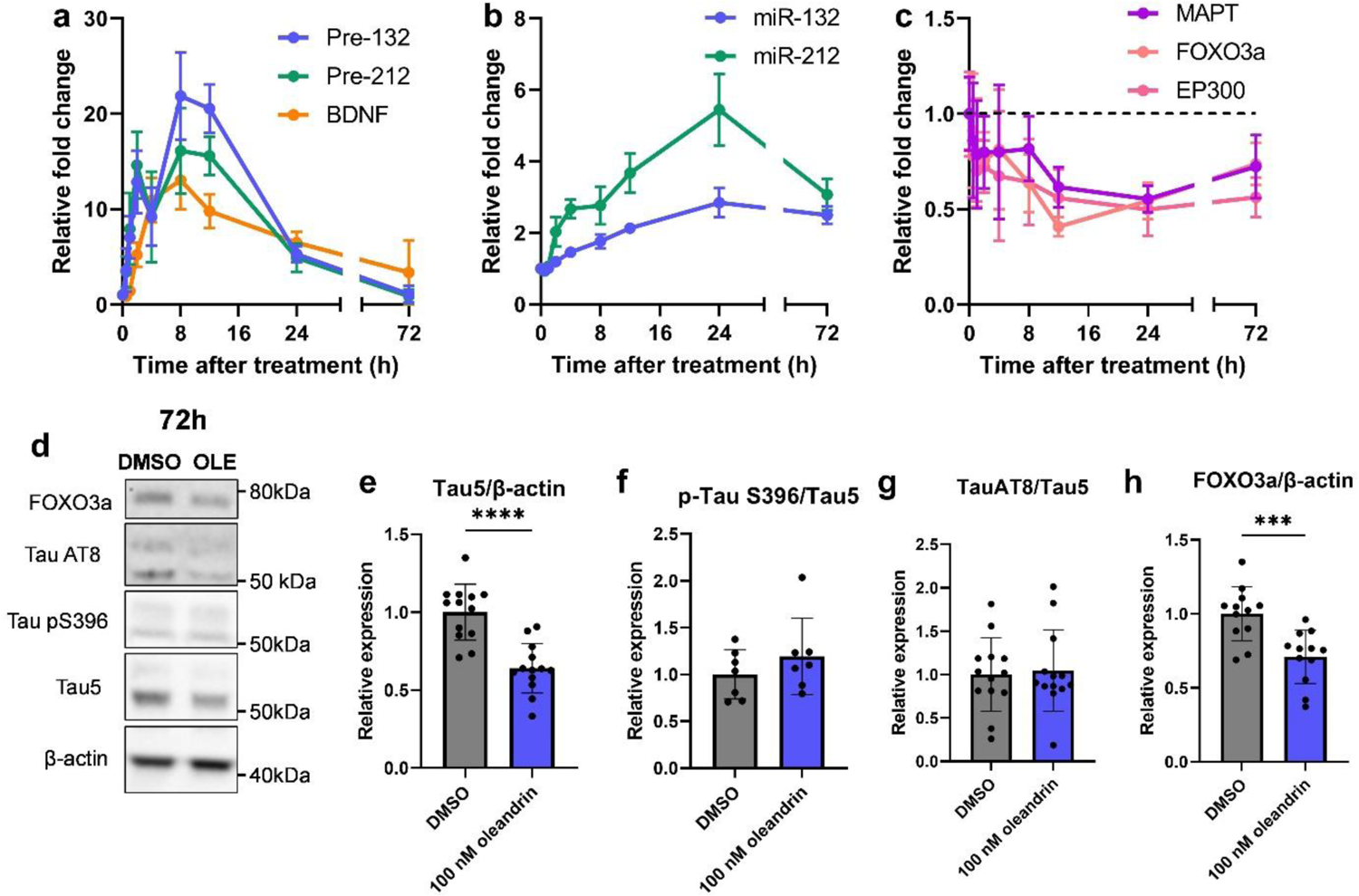
Oleandrin upregulated miR-132 and downregulated its targets over time. **a-c**, 100 nM oleandrin upregulated pre- and mature miR-132/212 and downregulated their mRNA targets over time (RT-qPCR analysis). **d-h**, Oleandrin downregulated total Tau, pTau (AT8 and S396), and FOXO3a protein after 72h treatment (Western blot analysis, unpaired two-tailed Student’s t-test, N= 4-8, error bars represent SD).

We hypothesized that the increase in miR-132 expression would lead to the downregulation of its targets. Indeed, we observed a time-dependent downregulation of *MAPT, FOXO3a, and EP300* mRNAs that matched the upregulation of miR-132 (Figure 4C). mRNA targets were significantly reduced to ∼50% of baseline at 24h and to ∼75% of baseline at 72h (Figure 4C), which was similar to the observed effects for miR-132 mimics 72h after transfection (S4A-C). Tau, pTau S202/T305 (AT8), pTau S396, and FOXO3a proteins were also downregulated, though the ratio of pTau: total Tau was unchanged (Figure 4D-H). In primary Tau wild-type (WT) and PS19 mouse neurons that overexpress human mutant Tau-P301S ^30^, 100 nM oleandrin upregulated miR-132 and downregulated both mouse *MAPT* and human *MAPT* after 72h treatment (S5D-G).

### Cardiac glycosides protected mature neurons against glutamate and Aβ toxicity

We hypothesized that the upregulation of miR-132 by the cardiac glycosides would be neuroprotective ^16^. As several studies have reported possible neurotoxic effects associated with cardiac glycosides ^31, 32^, we first treated rat neurons at different ages *in vitro* (DIVs 7/14/21/28) with digoxin, oleandrin, and proscillaridin A for 96h before measuring cellular viability (Figure 5A). Interestingly, DIV7 neurons were highly susceptible to cardiac glycoside toxicity, with significant loss of viability observed at the miR-132 EC_100_ for all compounds tested (Figure 5B-D). However, mature neurons were more resistant to cardiac glycoside toxicity, and no loss of viability was observed at miR-132 EC_100_ for neurons treated at DIV14 or later (Figure 5B-D).

**Figure 5.**
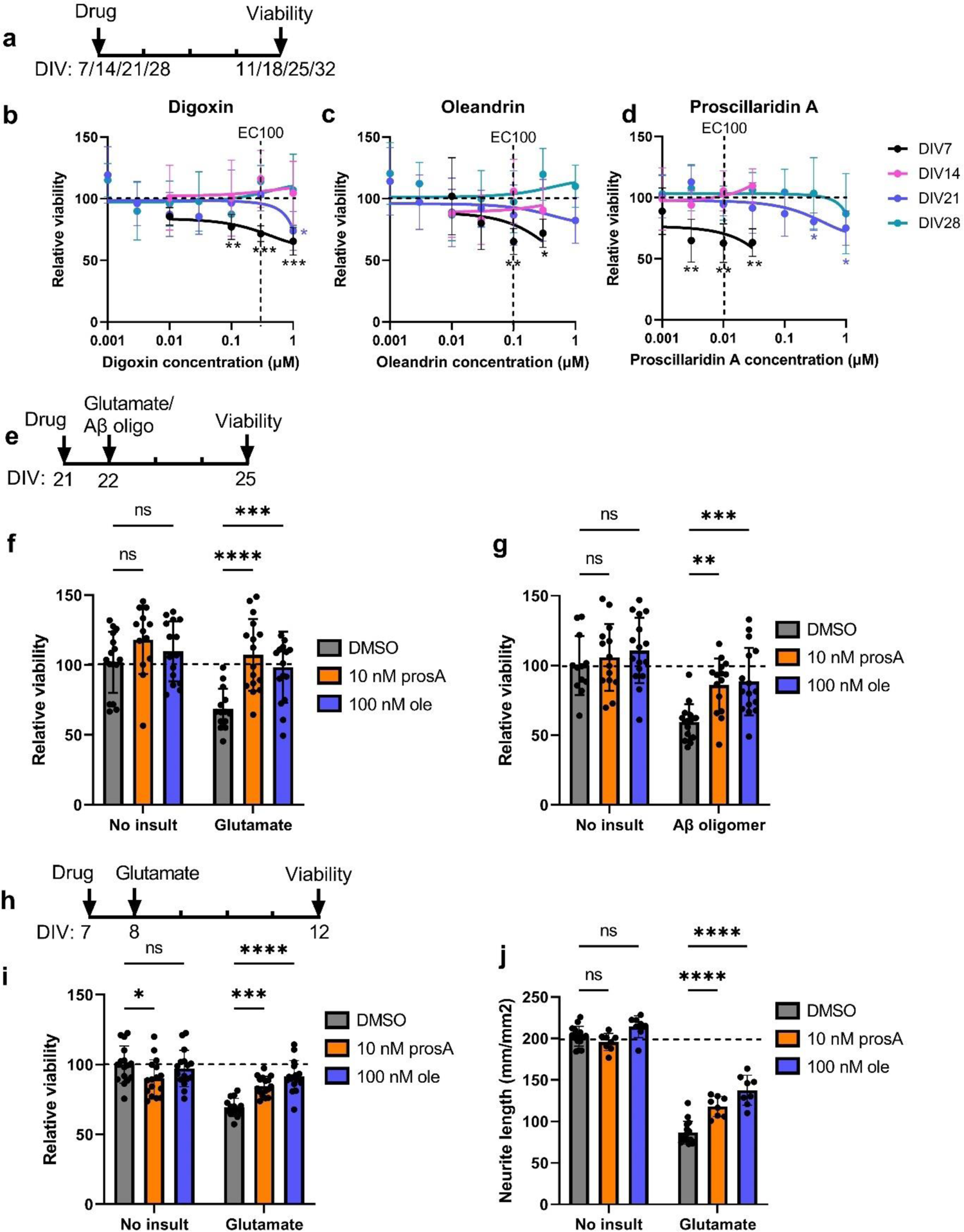
Cardiac glycosides rescued glutamate and Aβ oligomer-induced toxicity in primary rat neurons. **a-d**, Younger neurons were more susceptible to cardiac glycoside toxicity, whereas more mature neurons were resistant. Primary rat neurons were treated with various doses of digoxin, oleandrin, and proscillaridin A for 96h before viability was measured using WST-1. Cells treated at DIV7 showed a dose-dependent reduction in viability. In contrast, cells treated at DIV14, 21, or 28 showed little loss of viability, particularly at EC100 for miR-132 upregulation (unpaired t-test comparing to DMSO condition for each dose, N=4-8 per dose, error bars represent SD.). **e-g**, For DIV21 neurons, proscillaridin A and oleandrin were not toxic at baseline and fully rescued viability loss due to glutamate or Aβ oligomer treatment (2-way ANOVA, followed by Šídák’s multiple comparisons test, N=8-16 per condition, error bars represent SD). **h-j,** Proscillaridin A was mildly toxic to DIV7 neurons at baseline. However, both proscillaridin A and oleandrin fully rescued viability loss and partially rescued neurite loss due to glutamate treatment (2-way ANOVA, followed by Šídák’s multiple comparisons test, N=8-16 per condition, error bars represent SD).

To investigate neuroprotective effects, we first treated DIV21 rat neurons with oleandrin and proscillaridin A at EC_100_ for 24h, followed by 100 μM glutamate or 10 µM Aβ42 oligomers (Figure 5E). Pretreatment with proscillaridin A and oleandrin rescued neuronal viability loss due to toxic insults without affecting viability at baseline (Figure 5F, G). As we previously showed that miR-132 mimics rescued loss of viability in younger neurons treated with glutamate ^16^, we performed similar experiments in DIV7 neurons (Figure 5H). We observed a small loss of viability due to proscillaridin A at baseline (Figure 5I). However, both oleandrin and proscillaridin A rescued loss of viability caused by glutamate excitotoxicity (Figure 5I). Oleandrin and proscillaridin A also partially and dose-dependently rescued neurite loss induced by glutamate without affecting neurite at baseline (Figure 5J, S5).

### Cardiac glycosides significantly reduced Tau and pTau in human iPSC-neurons

To investigate the effects of cardiac glycosides in human neurons, we utilized two additional iPSC-derived neural progenitor cell (NPC) lines: MGH-2046-RC1 derived from an individual with FTD carrying the autosomal dominant mutation Tau-P301L (referred here as P301L), and MGH-2069-RC1 derived from a healthy individual directly related to MGH-2046 (referred here as WT). These NPCs, when differentiated into neurons (iPSC-neurons) for 6-8 weeks, represent well-established models for studying tauopathy phenotypes in patient-specific neuronal cells relative to a WT control ^33–35^.

Since Tau metabolism is regulated by miR-132 ^14^, and Tau lowering is a promising therapeutic strategy for ADRD ^36^, we first investigated the effects of cardiac glycosides on Tau protein levels. All three tested cardiac glycosides strongly and dose-dependently downregulated Tau, as exemplified by proscillaridin A. The treatment led to a clear reduction in total Tau (TAU5 antibody) and pTau S396 in WT neurons (Figure 6A) and in P301L mutant neurons after 24h and 72h (Figure 6D). For total Tau (TAU5), the upper band (>50 kDa, monomeric Tau + post-translational modifications (PTMs)) was more intense at lower drug concentrations. With increasing concentrations, the upper band disappeared, whereas the lower band (<50kDa, possibly non-pTau) became slightly more intense. This downward band shift suggested that proscillaridin A reduced both Tau accumulation and altered PTMs. Consistent with the latter, proscillaridin A reduced the monomeric form of pTau S396 (∼50 kDa) as well as the high molecular weight oligomeric pTau (≥250 kDa, Figure 6D).

**Figure 6.**
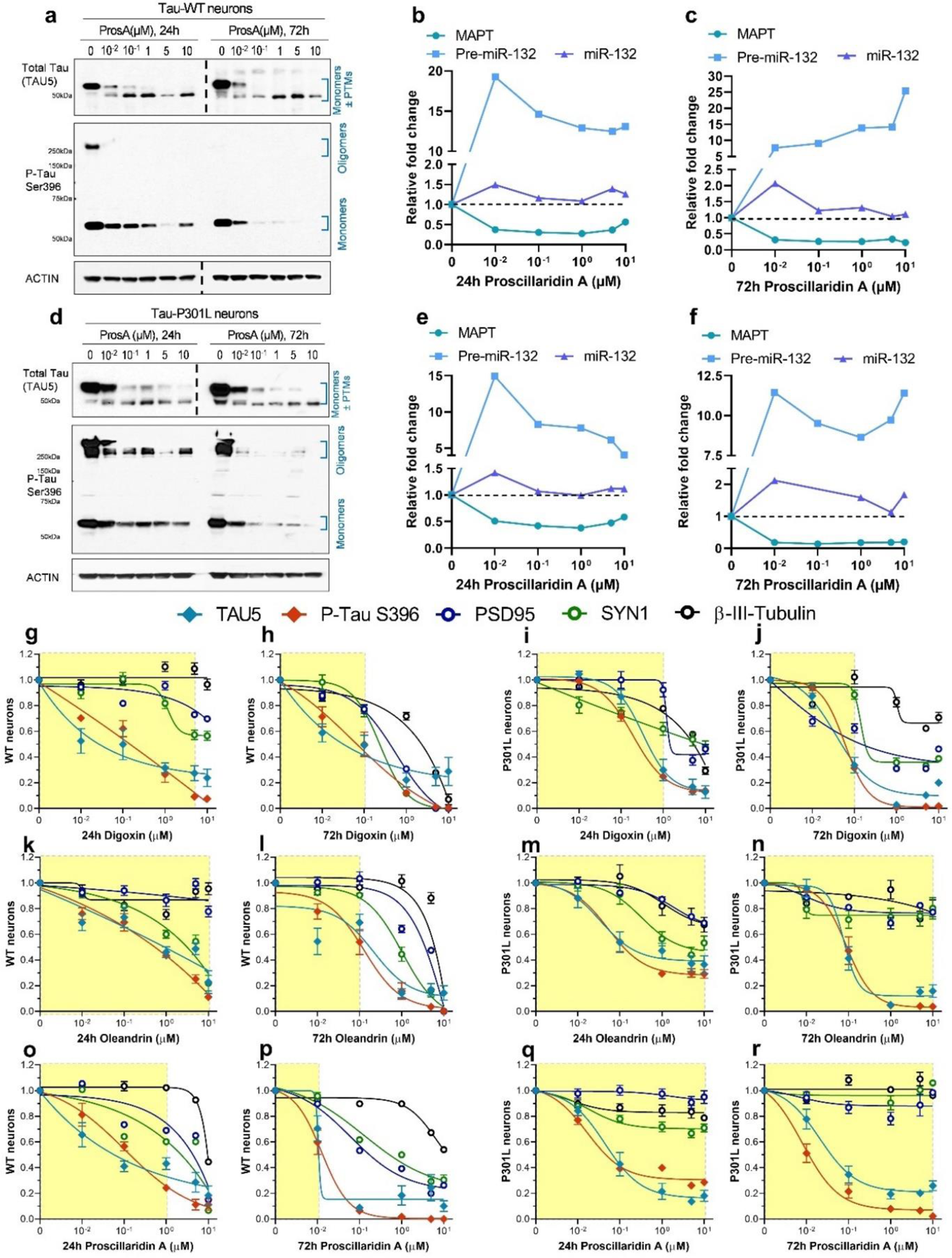
Dose-dependent reduction in Tau in human iPSC-neurons treated with cardiac glycosides. WT and P301L neurons were differentiated for 6 weeks, then treated with cardiac glycosides for 24h or 72h. **a**, Representative western blot for WT neurons treated with proscillaridin A (ProsA). A dose-dependent reduction in total Tau and p-Tau S396 was observed at both 24h and 72h. **b-c**, In parallel, a reduction in MAPT mRNA and an increase in pre-miR-132 and miR-132 RNA were observed. **d-f**, Similar results were also observed in Tau P301L neurons by western blot (d) and mRNA (e, f) analysis. **g-r**, Western blot densitometry quantification of dose-dependent effects on Tau (TAU5), pTau S396, and the synaptic makers PSD95 and SYN1 in WT and P301L neurons treated for 24h or 72h. The yellow shades indicate compound concentrations leading to <30% loss of at least two synaptic/microtubule markers (N=1-2, error bars represent SEM, the dotted lines indicated that separate Western blots were put together).

RT-qPCR was performed on a matched set of WT and P301L iPSC-neurons and showed a dose-dependent reduction in MAPT mRNA, a large increase in pre-miR-132, and a more modest increase in mature miR-132 (Figure 6 B, C, E, F). Similar results were obtained with digoxin and oleandrin treatments (S6). Further immunoblot results showed that in P301L iPSC-neurons, 72 h treatment with 1 µM proscillaridin A, digoxin, or oleandrin treatment reduced both soluble and insoluble total Tau and pTau S396 (S7A-C). 72h treatment also resulted in a dose-dependent reduction in miR-132 targets at the protein levels, including FOXO3a, EP300, GSK3β, and RBFOX1 (S7D-O).

For all compounds, the concentration of 10 µM was associated with >70% reduction in Tau and pTau with 24h and 72h treatments. However, this concentration also reduced neuronal synaptic markers, including post-synaptic density protein 95 (PSD95), synapsin 1 (SYN1), and β-III-tubulin representative of microtubules’ structural integrity. These results suggest that at high concentrations and with prolonged exposure, cardiac glycosides can compromise neuronal integrity. Nevertheless, for each drug, we observed a significant safety window in which Tau lowering was not associated with reduced synaptic or microtubule markers (Figure 6G-R). In all graphs, the yellow shade indicates the dose range where the loss of at least 2 synaptic markers was 30% or less (Figure 6G-R). Interestingly, WT neurons appeared more susceptible to loss of synaptic markers upon treatment than P301L neurons, particularly at 72h. For example, proscillaridin A was much less toxic to P301L neurons than WT neurons (Figure 6O, P, Q, R).

### Cardiac glycosides were neuroprotective in NPC-derived neuronal models of tauopathy

To examine the effects of the cardiac glycosides on neuronal viability, WT and P301L iPSC-neurons were treated with various doses of digoxin, oleandrin, and proscillaridin A for 24h or 72h. A dose-dependent loss of viability was observed with all three compounds, particularly at 72h. In Tau-WT neurons, there was up to 30% loss of viability after 72h treatment, particularly at the highest dose of 10 μM (Figure 7A-C). Interestingly, in Tau-P301L neurons, the toxicity observed was minimal, with <10% viability loss at the highest concentrations at 72h (Figure 7 D-F). These results were consistent with the previous immunoblot data (Figure 6G-R), showing that P301L neurons were more resistant to cardiac glycoside toxicity than WT neurons.

**Figure 7.**
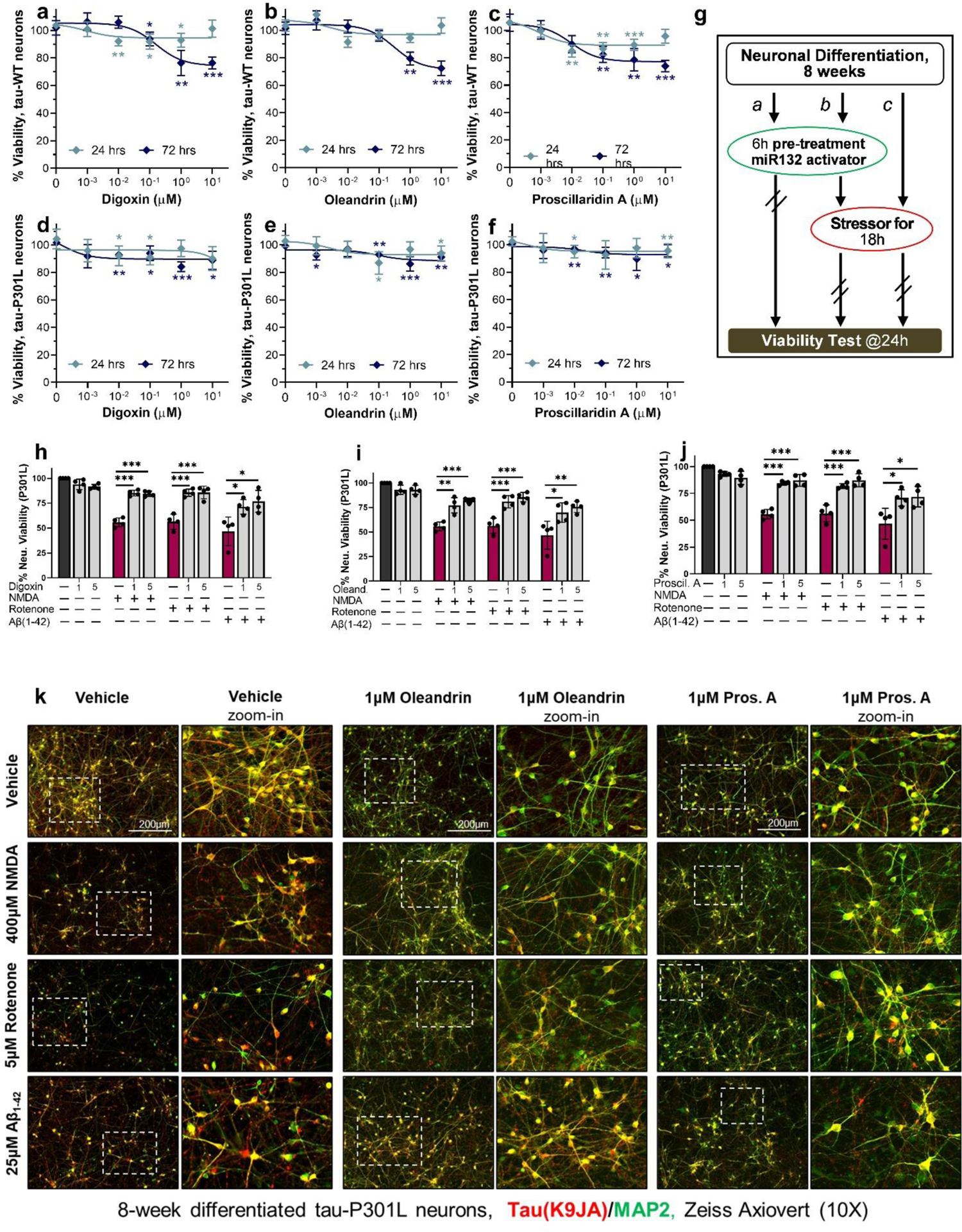
Cardiac glycosides rescued Tau-P301L neuronal vulnerability to stress. **a-f**, Compounds concentration effect on neuronal viability after 24h or 72h treatment of WT (a-c) and P301L (d-f) neurons. Data points indicate mean ±SD (N=2); unpaired two-tailed Student’s t-test. **g**, Schematic of the assay used to measure neuroprotective effects by cardiac glycosides in tauopathy neurons. **h-j**, Cardiac glycosides rescued the loss of viability in P301L neurons due to NMDA, rotenone, or Aβ42 oligomer treatment. Graph bars and data points show mean values ±SEM (N=2); unpaired two-tailed Student’s t-test. **k**, Representative images for P301L neurons at 8 weeks of differentiation treated with cardiac glycosides and each stressor compound. Total Tau (K9JA antibody) staining is shown in red, and MAP2 is shown in green. Scale bars are 200 μm.

We next tested whether cardiac glycosides can protect human neurons from various cell stressors that specifically affect human iPSC-neurons expressing mutant Tau ^35^. These include the excitotoxic agonist of glutamatergic receptors NMDA, an inhibitor of the mitochondrial electron transport chain complex I, rotenone, and the aggregation-prone Aβ (1–42) amyloid peptide. P301L neurons differentiated for 8 weeks were pretreated with cardiac glycosides for 6h prior to the addition of stressors for 18h, and viability was measured at the 24h time point (Fig.7g). Cardiac glycosides were added at the concentrations of 1 μM and 5 μM, which did not affect cell viability in P301L neurons at 24h (Figure 7 D-F). All cardiac glycosides significantly rescued neuronal viability in the presence of stressors (Figure 7H-J). The rescue could also be observed with immunofluorescent staining (Figure 7K). At baseline, 1 μM of digoxin, oleandrin, or proscillaridin A reduced Tau staining in agreement with the immunoblot data (Figure 6D) without visibly affecting neuronal health. Treatment with the stressors led to a significant loss of neurites and cell bodies in neurons pretreated with vehicle alone, which was rescued by pretreatment with the cardiac glycosides. Overall, these results demonstrate that low concentrations of cardiac glycosides were neuroprotective in human tauopathy neurons.

### Transcriptome analysis of human iPSC-neurons confirmed shared pathways affected by cardiac glycosides

To uncover the molecular effects of cardiac glycosides on neuroprotection beyond miR-132 and its canonical targets, we profiled transcriptomes of human iPSC-neurons after 72h of treatment with increasing doses of digoxin, oleandrin, proscillaridin A or vehicle alone (0.1% DMSO) using RNA sequencing (Figure 8A). Starting from low doses, cardiac glycosides remarkably changed the global transcriptome of P301L neurons as seen in principal component analysis (PCA, Figure 8B), with single principal component (PC1) being able to clearly separate controls from treatments. More importantly, three different cardiac glycosides regulated transcriptomes similarly and in a prominent dose-dependent manner (Figure 8B). Differential expression analysis identified thousands of genes significantly regulated with fold-change higher than 4, even though the statistical power was weakened by the intrinsic variance of dosage gradient (Figure 8C). Many genes were related to neuronal health and activity, including the strongly upregulated *ARC* which encapsulates RNA to mediate various forms of synaptic plasticity ^37, 38^, and downregulated *MAPT* and the *SLITRK3/4/6* family which plays a role in suppressing neurite outgrowth ^39^. We further focused on the biological pathways that were commonly regulated by all three cardiac glycoside compounds. Notably, these treatments affected a substantial number of shared pathways (Figure 8D). Many downregulated genes belong to 74 pathways related to neuronal development, morphology, health, or activity (Figure 8E). Upregulated genes were highly enriched in positive regulators of transcription, negative regulators of programmed cell death, and regulators of stress and the unfolded protein response (Figure 8F). Furthermore, dozens of transcription factors that had binding sites on MIR132 promoter and may upregulate its expression, including *CREB5*, were commonly upregulated by cardiac glycosides (Figure 8G). The neuroprotective *BDNF* signaling pathway was significantly upregulated (S8), corroborating our previous observation that cardiac glycosides upregulated B*DNF* in rat neurons (Figure 4A). Therefore, while digoxin, oleandrin, and proscillaridin A all induced miR-132 expression, they likely also regulated multiple other pathways. Overall, shared transcriptomic alterations and regulated pathways further confirmed the common molecular mechanisms of action of cardiac glycosides and their ability to activate stress-protective programs in highly vulnerable Tau-mutant neurons (Figure 8H).

**Figure 8.**
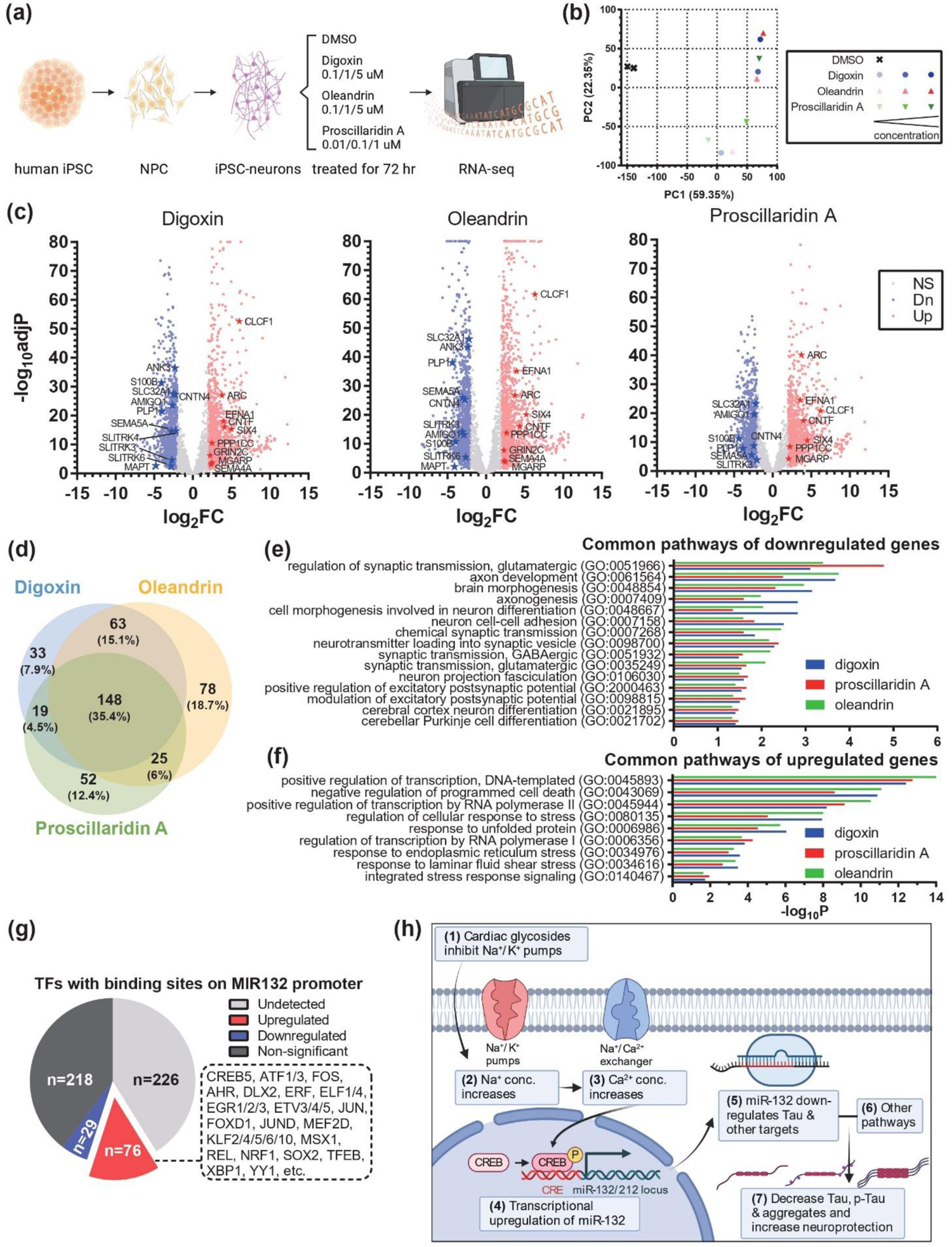
Transcriptome analysis of human iPSC-neurons treated with cardiac glycosides. **a**, Workflow of the experiment design. **b**, Principal component analysis (PCA) indicated the strong and dose-dependent alteration of global transcriptomic profiles after treatments. **c**, Volcano plots showed significant down- and up-regulated genes, labeled in blue and red dots, respectively. Stars highlighted dysregulated genes involved in neuronal activity and health. **d**, Venn diagram indicated the similarity of pathways affected by three cardiac glycosides. **e**, Selected neuronal pathways highlighted in common pathways of down-regulated genes. **f**, Selected transcription- and response-related pathways highlighted in common pathways of up-regulated genes. **g**, Effects of cardiac glycosides on the expression of transcription factors (TFs) that have binding sites on MIR132 promoter. **h**, Working model showing the effects of cardiac glycosides: cardiac glycosides act through their conventional mechanism leading to the transcriptional upregulation of miR-132. The increase in miR-132, together with other pathways altered by cardiac glycosides, downregulated various forms of Tau and provided neuroprotection against toxic insults.

## DISCUSSION

As miRNAs have been increasingly recognized as master regulators of many biological processes and promising therapeutic targets, screens for miRNA modulators have recently emerged. Several studies have reported successful screens for small molecules that inhibit the activity of oncogenic or pathogenic miRNAs, including miR-21 ^40, 41^, miR-122 ^42^, and miR-96 ^43^. Small-molecule inhibitors of miRNAs can be chemically modified to improve pharmacological properties and efficient CNS delivery, even though with potentially inferior target specificity relative to miRNA antisense oligonucleotides. On the other hand, miRNA mimic oligonucleotides require chemical modifications for stabilization and durable activity *in vivo*, which may reduce overall potency in the simultaneous regulation of multiple downstream targets. Therefore, small-molecule inducers of specific miRNAs could provide additional advantages as therapeutics. To date, no miRNA inhibitor or mimic oligonucleotide therapeutics have been FDA-approved, very few reporter-based screens have been published, and no systematic screens relying on broader miRNome-level readouts have been performed for small-molecule miRNA modulators ^22^.

Most HTSs for modulators of gene expression employ cell lines as screening platforms and gene-specific heterologous reporter systems as primary assays. However, proliferating, immortalized cells have limited value for identifying neuroprotective agents, and neurons are known to be technically difficult to transfect efficiently and uniformly, especially on a large scale ^21^. Here, we applied HTS with miRNA-seq to directly quantify expression levels of hundreds of miRNAs in human neurons treated with small molecule compounds. Notably, the present study is the first HTS-HTS for small RNAs that was enabled by the low-input requirement of RealSeq technology ^24^, although HTS-HTS for mRNA has been conducted previously ^44–46^. Despite the relatively small scale of ∼1900 compounds, we successfully validated 4 different classes of drugs that upregulate miR-132, most notably the cardiac glycosides family. As the first small molecule screen for neuronal miRNA modulators, the obtained dataset can be reanalyzed to identify compounds that regulate any of the ∼450 miRNAs, providing a unique new resource (Table S4) and facilitating further discoveries of miRNA-targeting drugs.

In this study, we focus on miR-132, a master neuroprotector. Several members of the cardiac glycoside family, Na+/K+ ATPase pump inhibitors, were successfully validated to upregulate miR-132/212 consistently. Of note, cardiac glycosides such as digoxin and digitoxin are widely used for treating congestive heart failure and cardiac arrhythmias. However, they have a narrow therapeutic index and can be toxic at high doses^47^. Recent studies have reported that cardiac glycosides are neuroprotective in animal models at low (sub-μM to μM) concentrations in stroke^48, 49^, traumatic brain injury ^50^, systemic inflammation ^51^, and AD and tauopathies ^52, 53^. Furthermore, clinical studies suggest that treatment with digoxin might improve cognition in older patients with or without heart failure ^54^. Our data supported that the cardiac glycosides reduced Tau accumulation and rescued Tau-mediated toxicity. Further work remains to be done to investigate if any member of the cardiac glycosides can be developed into effective and safe therapeutics for long-term treatment against neurodegenerative diseases. Oleandrin, which was previously shown to be neuroprotective with excellent brain penetrance and retention ^49, 55^, and proscillaridin A, which exhibited the lowest EC_50_ in rat and human neurons, may be good starting points. Furthermore, as the expression of ATP1A3 is restricted to neurons, whereas ATP1A1 and ATP1A2 are more ubiquitously expressed ^56^, drugs that selectively target the ATP1A3 isoform may alleviate the systemic impact of the cardiac glycosides such as on the cardiac system.

Several questions that emerge from our observations are worth investigating further. First, there is a significant difference in the fold change of mature and pre-miR132. Pre-miR-132 was upregulated by 10 to 30-fold, whereas mature miR-132 in the same treatment group was upregulated by only 1.5-3-fold (Figures 4, 6). We hypothesize that there may be physiological mechanisms that maintain the levels of mature miR-132 within a 2-fold difference, perhaps a bottleneck in processing pre-miR-132 to mature miR-132. Interestingly, miR-132 is downregulated by ∼1.5-2.5-fold in various neurodegenerative diseases ^7^, suggesting that the increase promoted by treatment with cardiac glycosides is sufficient to restore physiological miR-132 levels. Second, the effect of cardiac glycosides on downregulating human *MAPT* mRNA and Tau protein appears to be much stronger than for rodent Tau. After 72h treatment, 100 nM oleandrin downregulated rat *MAPT* mRNA by 27% and rat Tau protein by 35% (Fig. 4). The same treatment downregulated human *MAPT* mRNA by 87% and human Tau protein by 59% in mutant P301L neurons (Figure 6 and S6). Some differences may be attributed to differences in sensitivity to cardiac glycosides between rodents and humans, as mouse ATP1A1 is inhibited by ouabain and digitoxin at >100-fold higher concentration than human ATP1A1^57^. However, in PS19 mouse neurons, which express ∼8-fold more human *MAPT* than endogenous mouse *MAPT* mRNA, oleandrin downregulated both mouse and human *MAPT* by ∼40-50% (S4), suggesting that cardiac glycosides and miR-132 may target human *MAPT* more effectively. Third, several studies have proposed that cardiac glycosides downregulate *MAPT* and Tau and provide neuroprotection through other pathways, including increased autophagy ^52^, alternative splicing of *MAPT* mRNA^58^, increased BDNF ^28^, and inhibiting reactive astrocytes ^53^. Our transcriptomic results support that many neuronal pathways are altered, suggesting that cardiac glycosides can modulate multiple pathways that converge on the downregulation of Tau and increased neuroprotection. While further investigation is needed to determine the contribution of miR-132 upregulation to Tau downregulation and neuroprotection, cardiac glycosides emerge as promising therapeutics for neurological disorders, if they can be improved to reduce systemic toxicity and enhance brain penetrance and retention.

In summary, our pilot HTS-HTS of miRNA regulators on human neurons discovered the cardiac glycoside family as novel miR-132 inducers. These compounds specifically upregulated miR-132 expression via transcription activation by inhibiting the Na^+^/K^+^ ATPases and could protect rat primary neurons and a human iPSC-derived neuronal model of tauopathy against diverse insults, including glutamate, Aβ oligomers, NMDA, and rotenone. Our pilot study not only highlights cardiac glycosides as promising treatments for neurodegenerative diseases but also provides a key omics resource for future neuronal miRNA regulator discoveries.

## Acknowledgments

This work was supported by the R56 AG069127 and the Rainwater Foundation/ Tau Consortium grants to A.M.K. S.J.H., and M.C.S. were supported by Rainwater Foundation/ Tau Consortium funding. T.L.Y.P., C.R.M., and J.M.S.S. were supported by R01AG055909. The BWH iPSC NeuroHub provided support for NGN2-iNs related work. The ICCB-Longwood Screening Facility provided the compounds and instruments for performing the high-throughput drug treatment. The NeuroTechnology Studio at Brigham and Women’s Hospital provided IncuCyte instrument access and consultation on data acquisition and data analysis. Dr. Bradford Dickerson (MGH), Dr. James Gusella (MGH), Diane Lucente (MGH), and Dr. Bruce Miller (UCSF) are thanked for the generous sharing of patient cell lines. Dr. Michelle Arkin (UCSF), Dr. Erik Uhlmann (DFCI), and Dr. Evgeny Deforzh (BWH) are thanked for their helpful discussion, comments, and edits. Ramil Arora and Harini Saravanan are thanked for annotating the compounds, as shown in Table S1. Dr. Rachid El Fatimy is thanked for the preparation of Ab oligomers. PubChem Sketcher was used to prepare the chemical structures in Supp. Table BioRender was used in the preparation of Figures 1, 8, and S8.

## Contributions

A.M.K., Z.W., and L.D.N. conceived and designed the study. Z.W. and L.D.N. equally contributed as first authors. Z.W., L.D.N., M.C.S., S.B.S., R.R., C.R.M., J.M.S.S., and C.H. performed experiments for this study. T.L.Y.P., S.J.H., and A.M.K. provided the resources needed for experiments. L.D.N. wrote an original draft of the manuscript. A.M.K, Z.W., and L.D.N. reviewed and edited the manuscript. All authors reviewed and commented on the manuscript.

## Corresponding authors

Correspondence to Zhiyun Wei or Anna M. Krichevsky.

## Availability of data and materials

miRNA-sequencing and mRNA-sequencing data that support the findings of this study will be deposited into Sequence Read Archive with accession number to be determined. Contact corresponding authors for requests of materials and cell lines used in the manscript.

## SUPPLEMENTAL DATA LEGEND

**Supplemental Figure S1:**
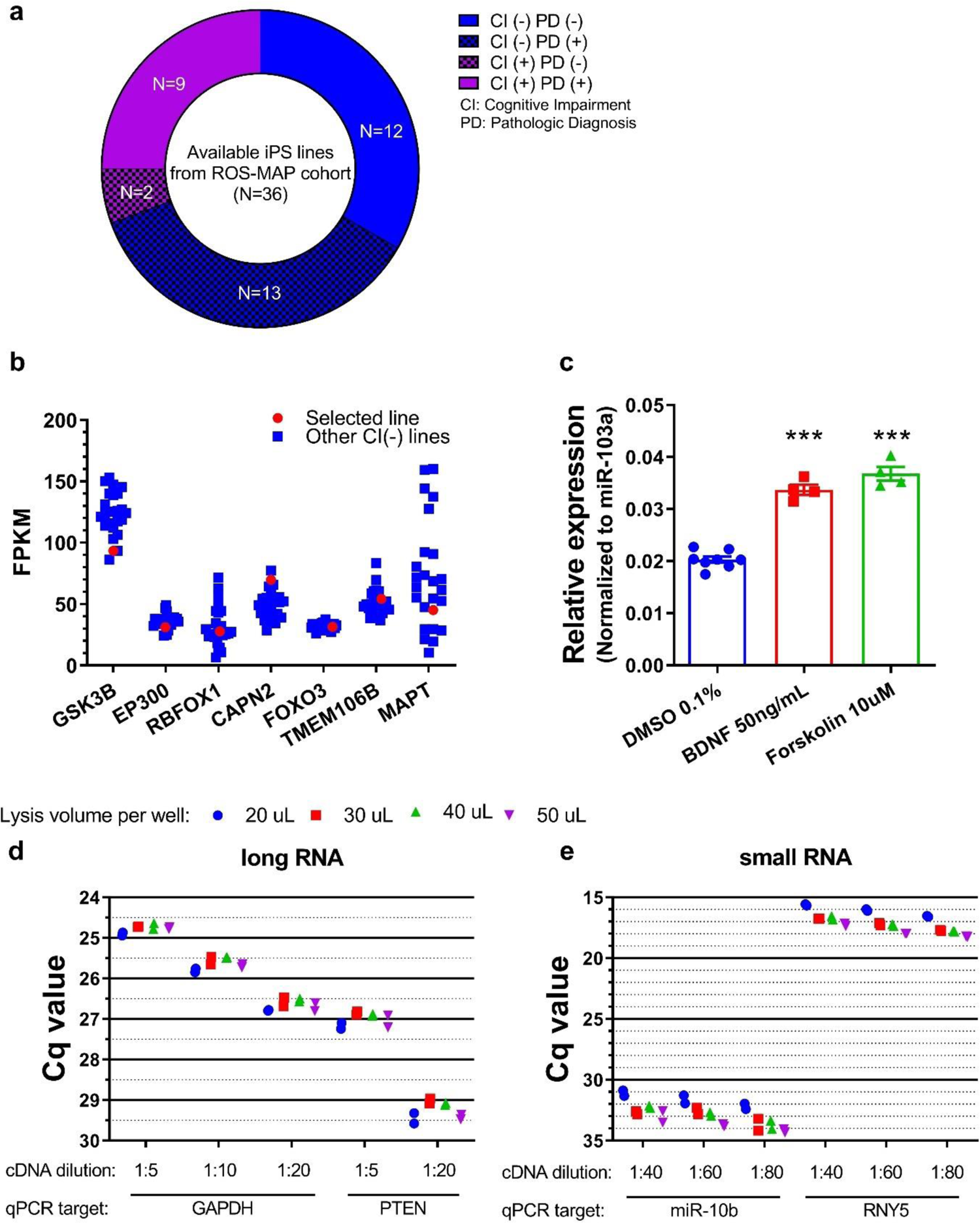
Optimization of screen setup. **a**, ROS-MAP NGN2-iN lines available. **b**, The BR43 line was selected among the NGN2-iNs established from donors without a clinical diagnosis of AD for its median expression of previously validated miR-132 targets. This line came from an 89-year-old female donor without clinical AD diagnosis but with pathological AD diagnosis. **c**, miR-132 was upregulated by positive controls forskolin and BNDF in BR43 NGN-2 iNs. **d-e**, 40 to 50 μL of Takara direct lysis buffer was optimal for direct lysing. 1:10 and 1:60 dilutions were optimal for qPCR on long RNA RT and small RNA RT, respectively.

**Supplemental Figure S2:**
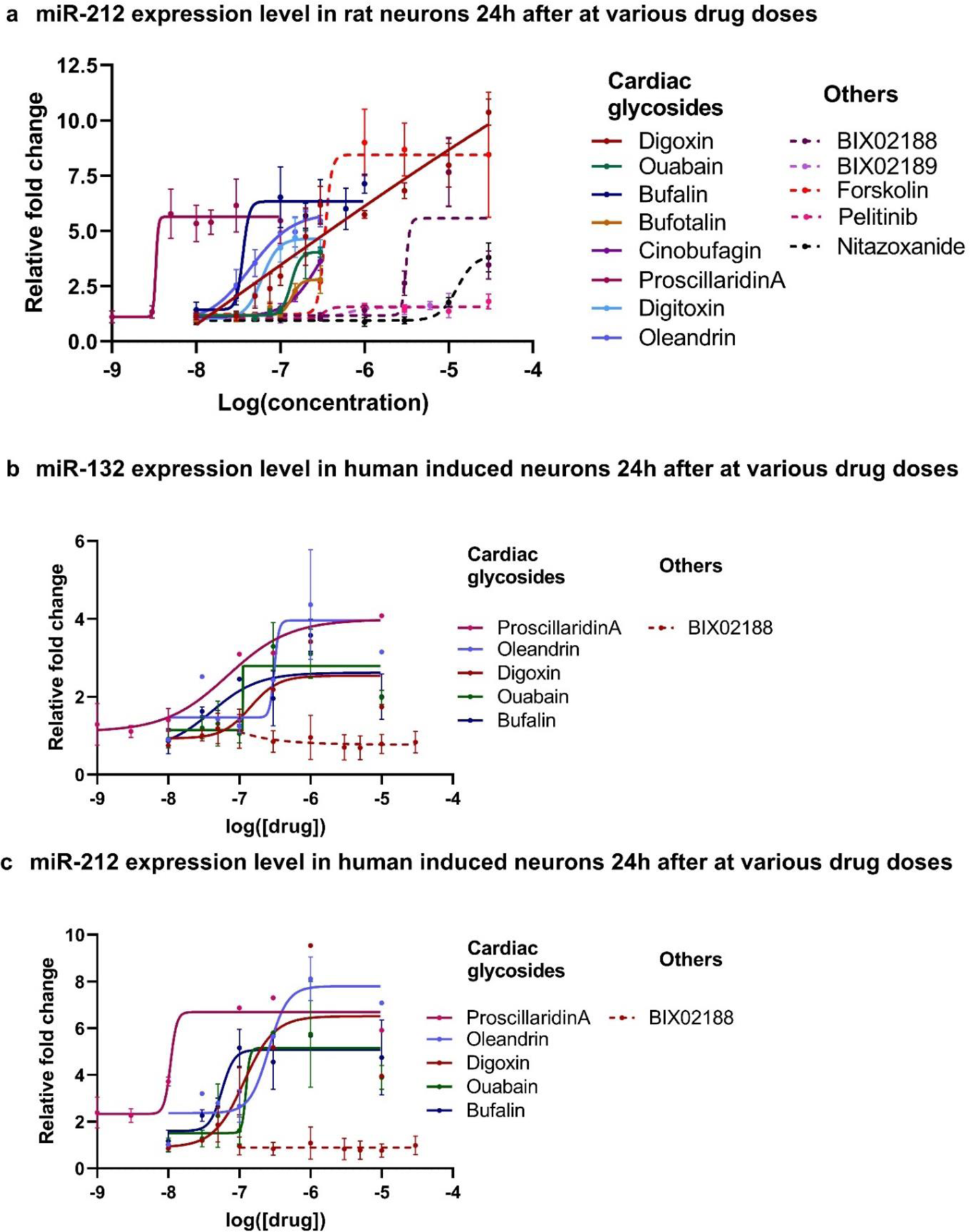
Dose-dependent upregulation of miR-132 and miR-212 in primary rat neurons and human NGN-2iNs. **a**, Dose curve experiments for miR-212 were performed in DIV14 rat neurons after 24h treatment (N=4-6). **b-c**, Dose curve experiments for miR-132 and miR-212 were performed in DIV21 human NGN2-iNs after 24h treatment (N=1-2). Solid lines were used for cardiac glycosides, and dotted lines were used for non-cardiac glycosides. EC_50_ and max fold change were calculated using sigmoidal fit, 4 parameters. Error bars represent SD.

**Supplemental Figure S3:**
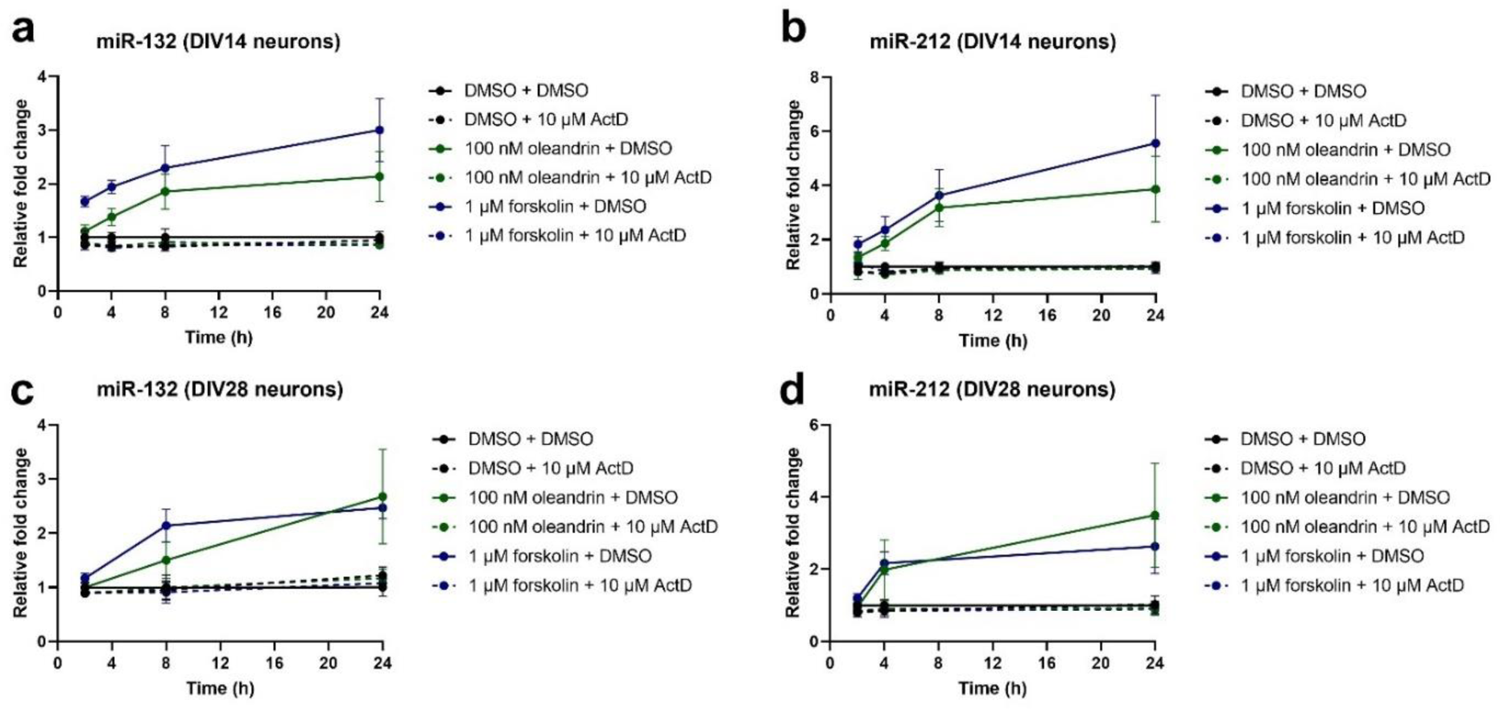
Actinomycin D inhibited the upregulation of miR-132 and miR-212. Time-dependent upregulation of miR-132 and miR-212 was completely abolished by pretreatment with 10 μM actinomycin-D before forskolin or oleandrin in DIV14 rat neurons (**a-b**) and DIV28 rat neurons (**c-d**).

**Supplemental Figure S4:**
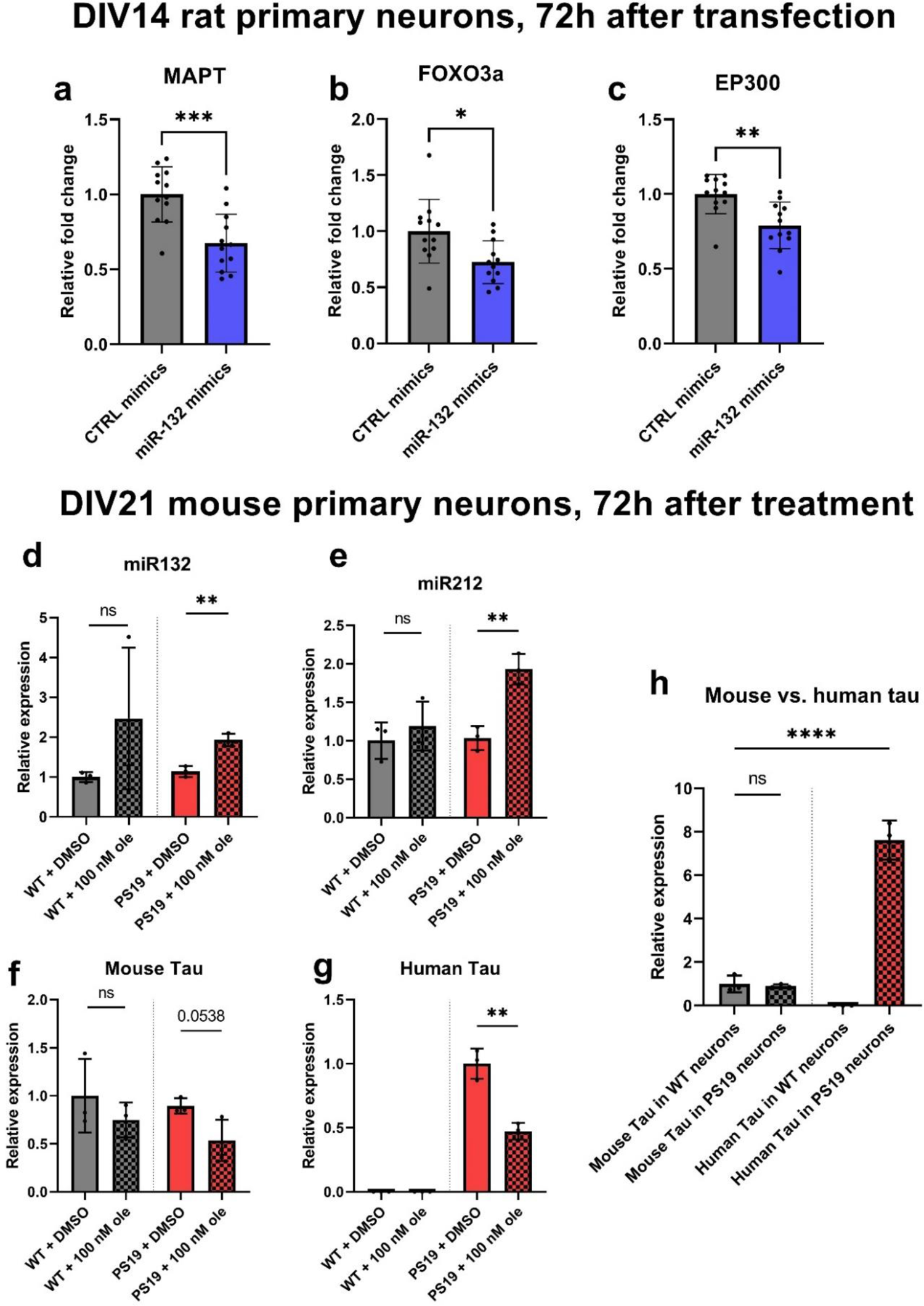
Additional effects of miR-132 mimics and cardiac glycosides. **a-c**, miR-132 target mRNAs were downregulated 72h after transfection with miR-132 mimics. **a**, MAPT. **b**, FOXO3a. **c**, EP300 (N=12, unpaired two-tailed Student’s t-test, Error bars represent SD). **d-g**, miR-132 and miR-212 were also upregulated and mouse and human MAPT mRNA were downregulated by oleandrin in primary PS19 mouse neurons. Similar observations were also observed in WT neurons but were not statistically significant. **h**, human MAPT mRNA was expressed at ∼8-fold higher than endogenous mouse MAPT (N=3, unpaired two-tailed Student’s t-test, error bars represent SD).

**Supplemental Figure S5:**
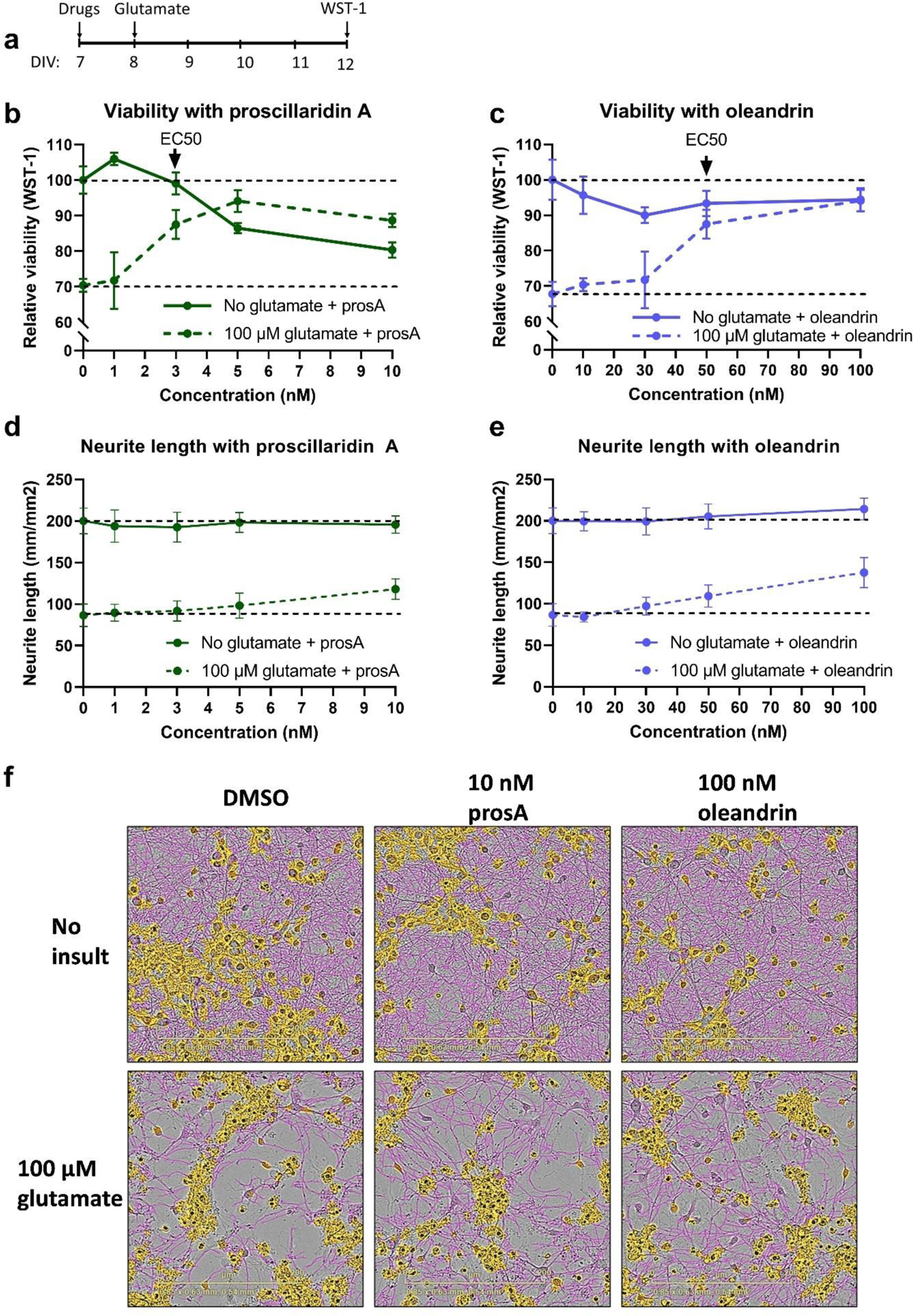
Oleandrin and proscillaridin A rescued viability loss from glutamate toxicity but were also mildly toxic in younger primary neurons. a, Experimental scheme. b, Proscillaridin A dose-dependently reduced baseline viability (solid line) but also dose-dependently rescued loss of viability due to glutamate (dotted line). c, Similar results were obtained for oleandrin. d-e, Proscillaridin A and oleandrin did not affect neurite length at baseline and partially rescued loss of neurites due to glutamate. f, Representative images of neurons treated with glutamate and proscillaridin A or oleandrin. Cell bodies were highlighted in yellow, and neurites were traced in pink. N=6-8, error bars represent SD. Scale bars are 200 µm.

**Supplemental Figure S6:**
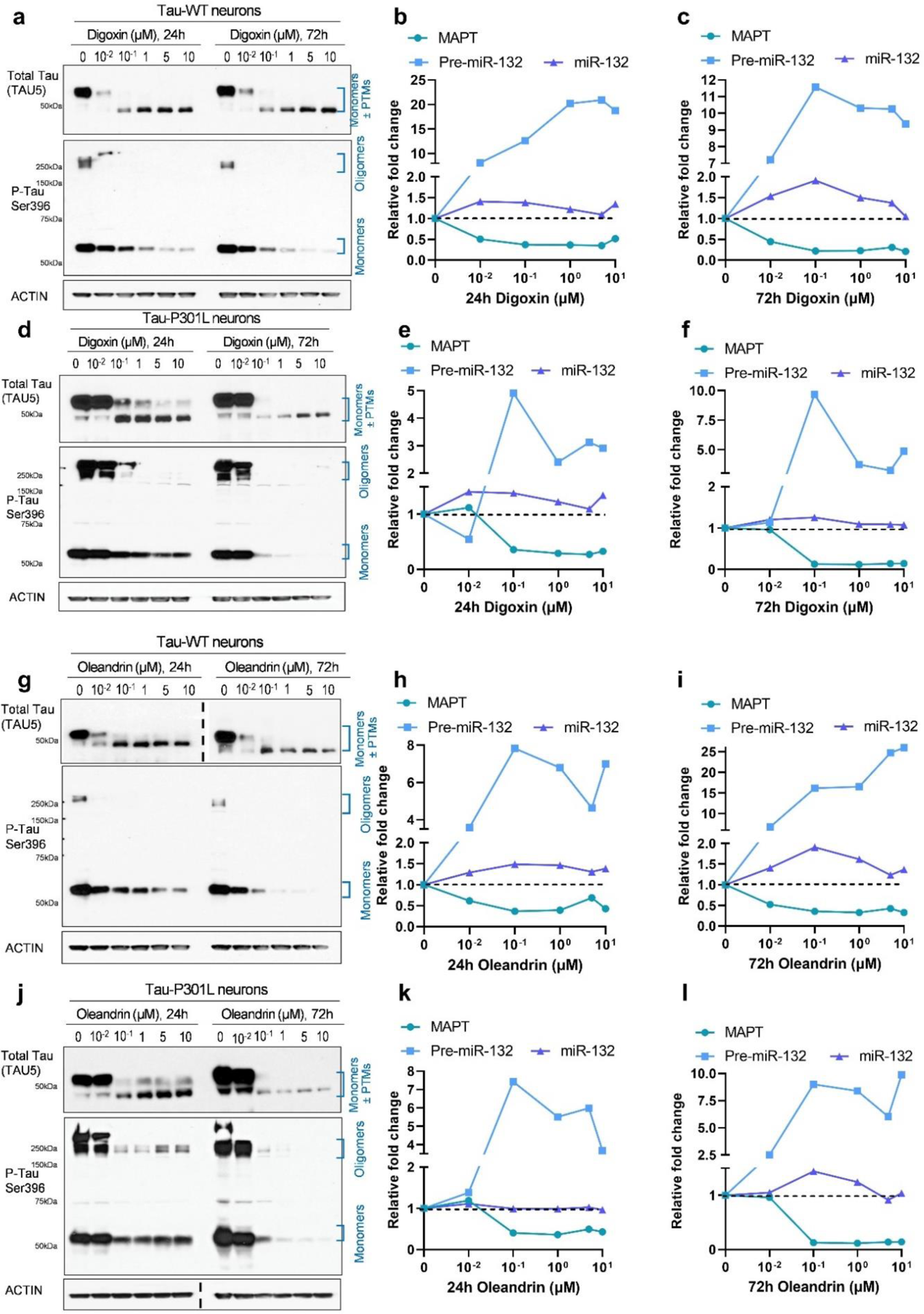
Dose-dependent reduction of Tau in iPSC-neurons treated with cardiac glycosides. Tau WT and P301L neurons were differentiated for 6 weeks, then treated with cardiac glycosides for 24h or 72h. **a**, Representative western blot for WT neurons treated with digoxin. A dose-dependent reduction in total Tau (TAU5) and p-Tau S396 was observed at both 24h and 72h. **b-c**, In parallel, a reduction in MAPT mRNA and an increase in pre-miR-132 RNA were observed. **d-f**, Similar results were also observed in Tau P301L neurons. **g-l,** Similar results were observed for WT and P301L neurons treated with oleandrin.

**Supplemental Figure S7:**
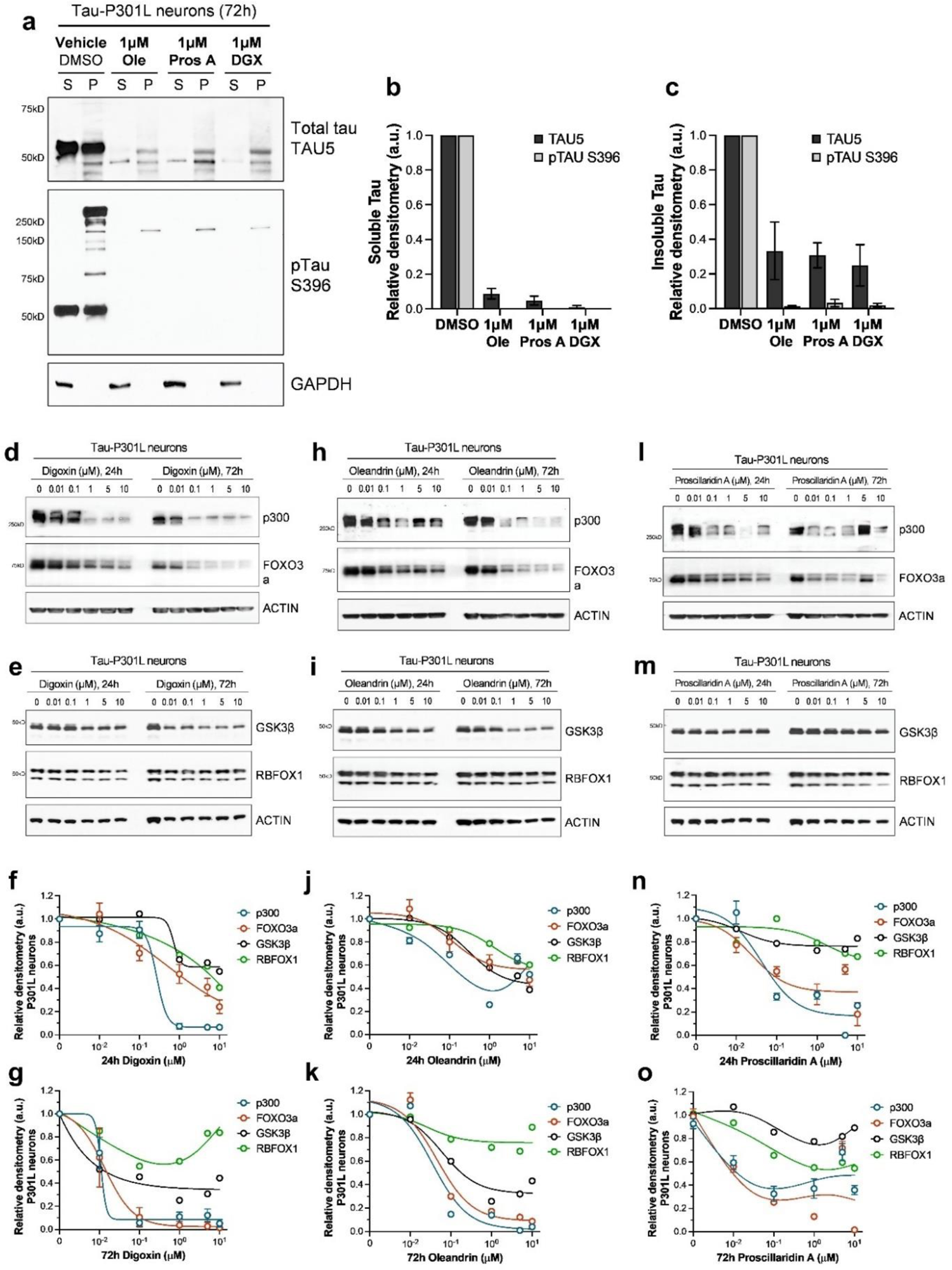
Cardiac glycosides’ effect on Tau solubility and miR-132 targets in P301L neurons. iPSC-neurons differentiated for 6 weeks were treated for 72h at concentrations of oleandrin (Ole), proscillaridin A (Pros A) and digoxin (DGX) leading to maximum Tau reduction without detectable toxicity. **a,** Representative western blot analysis of protein lysates generated by detergent fractionation for detection of total Tau (TAU5) and pTau S396 in the soluble (S) and insoluble-pellet (P) fractions. **b-c,** Densitometry analysis of the western blots (N=2). Graph bars represent mean densitometry ± SD for soluble (**b**) and insoluble (**c**) Tau levels relative to vehicle (DMSO). **d-g**, 24h and 72h treatment with digoxin resulted in a dose-dependent reduction in miR-132 targets, including p300, FOXO3a, GSK3β, and RBFOX1 (N =2). Error bars represent SEM. Similar results were obtained for oleandrin (**h-k**), and proscillaridin A (**l-o**).

**Supplemental Figure S8:**
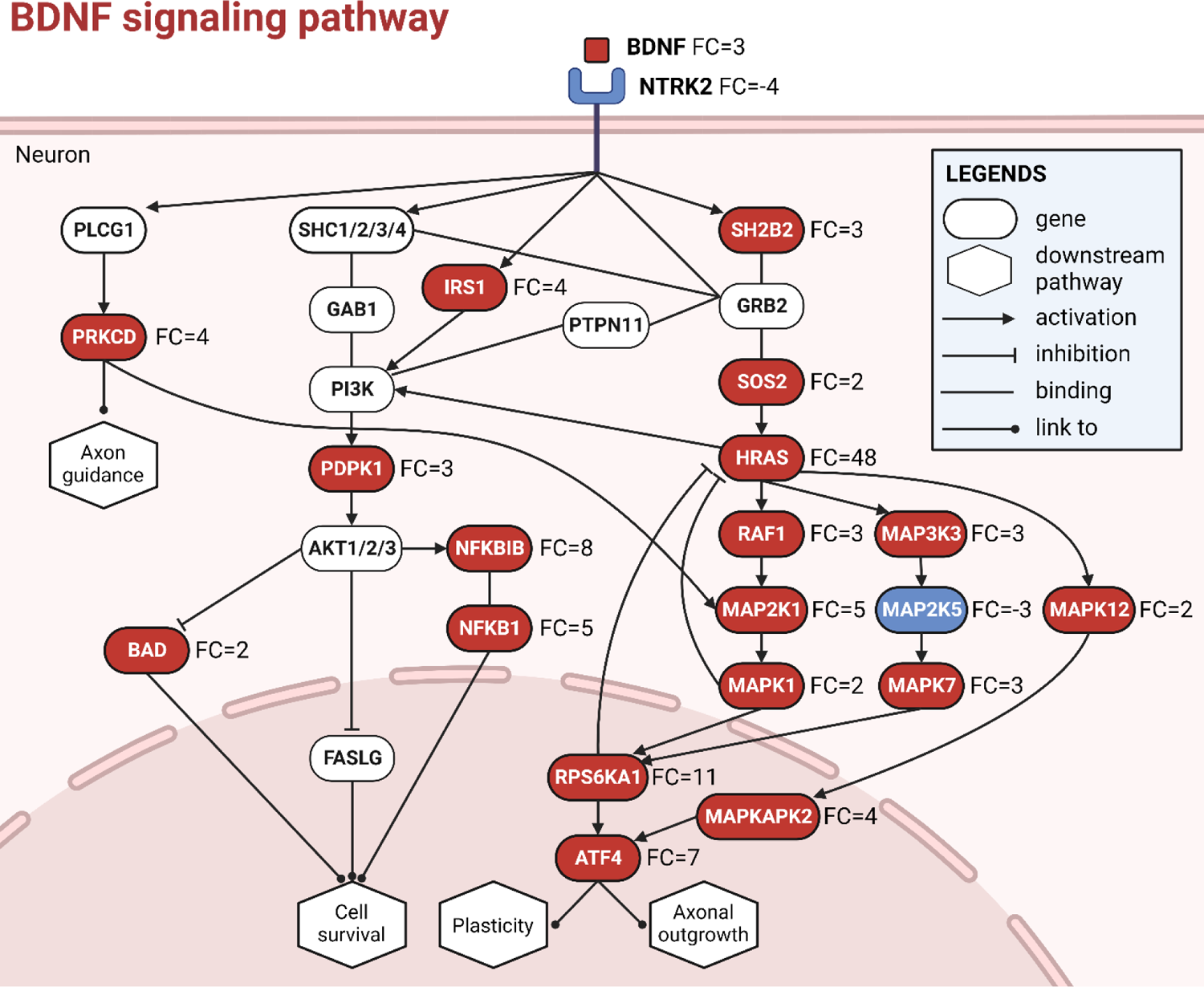
BDNF signaling pathway was enriched with genes upregulated by cardiac glycosides in iPSC-neurons. Pathway plot was modified based on KEGG neurotrophin signaling pathway. Genes in red, blue, and white boxes represented upregulated, downregulated, and unaffected ones, respectively. Mean fold changes (FC) among three cardiac glycosides were labeled, of which positive value represented upregulation and vice versa.

**Supplemental Table 1:**
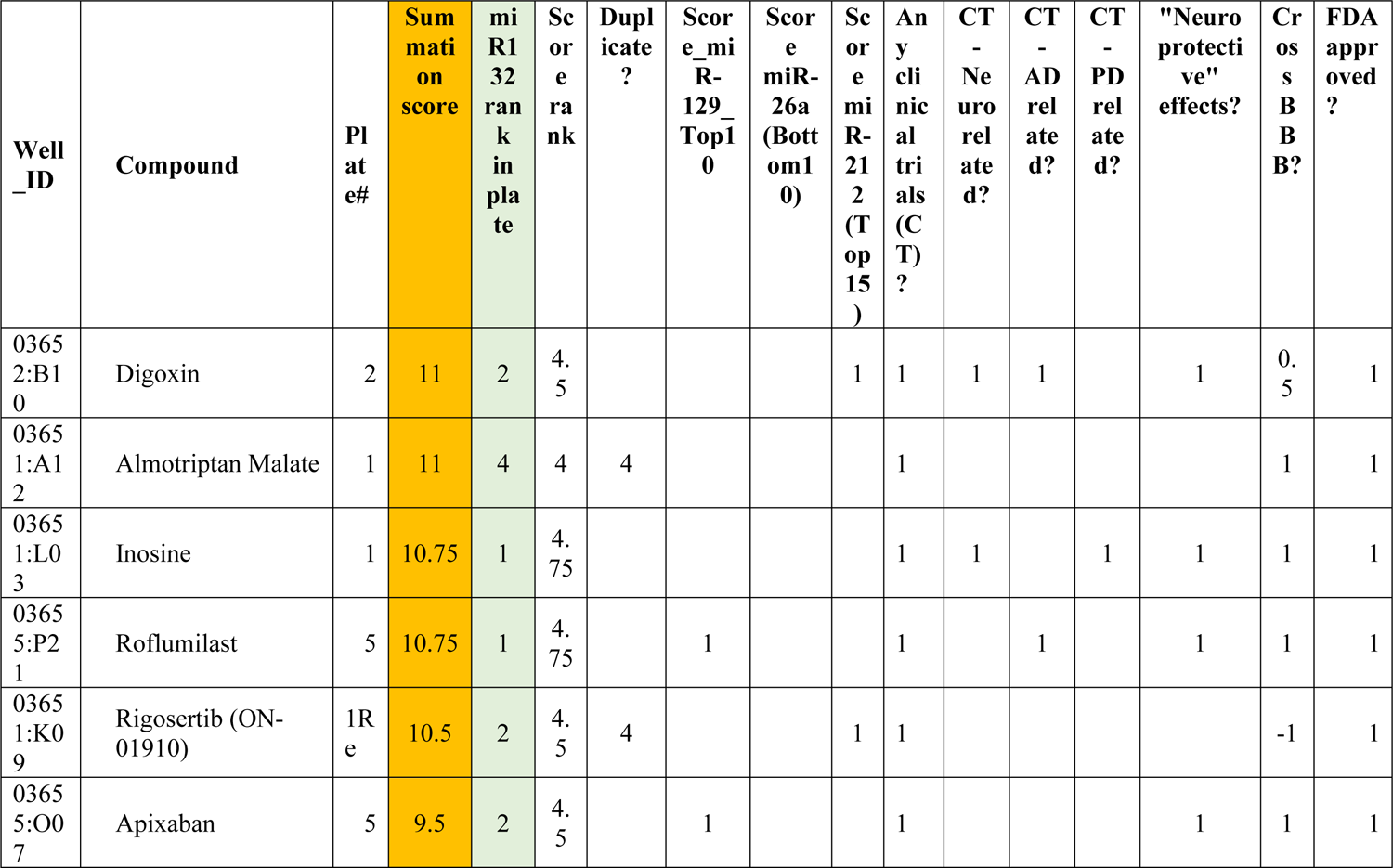
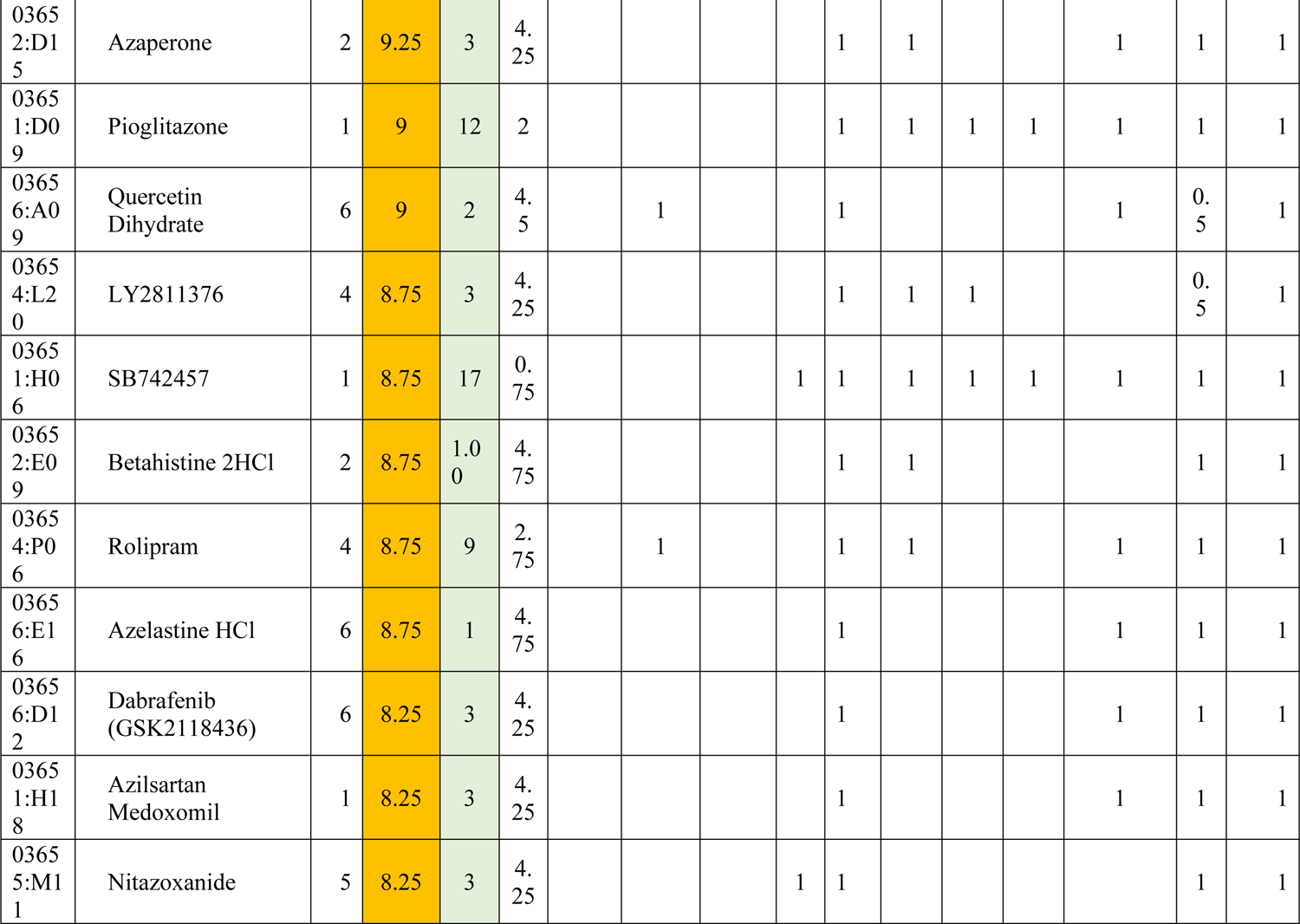
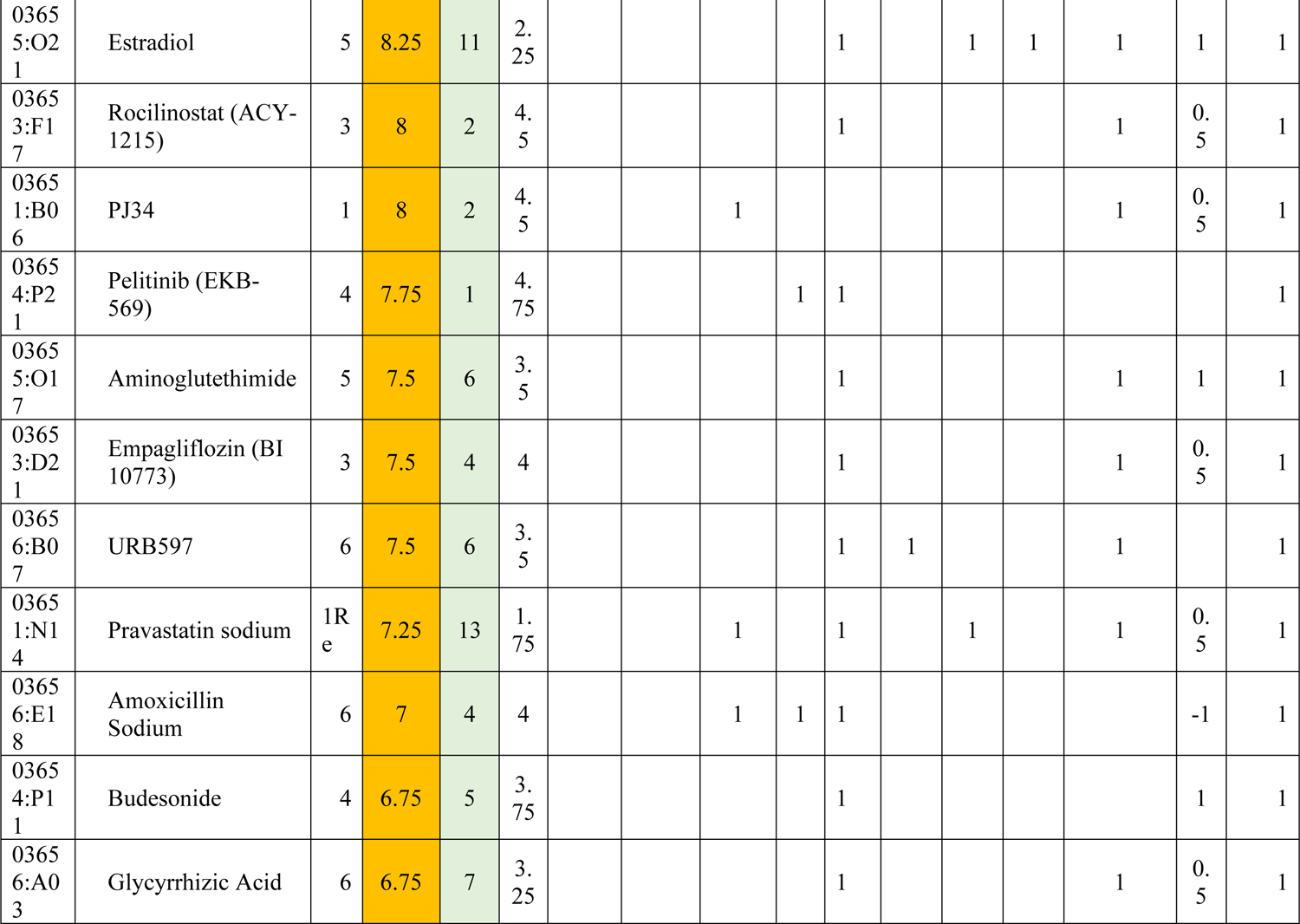
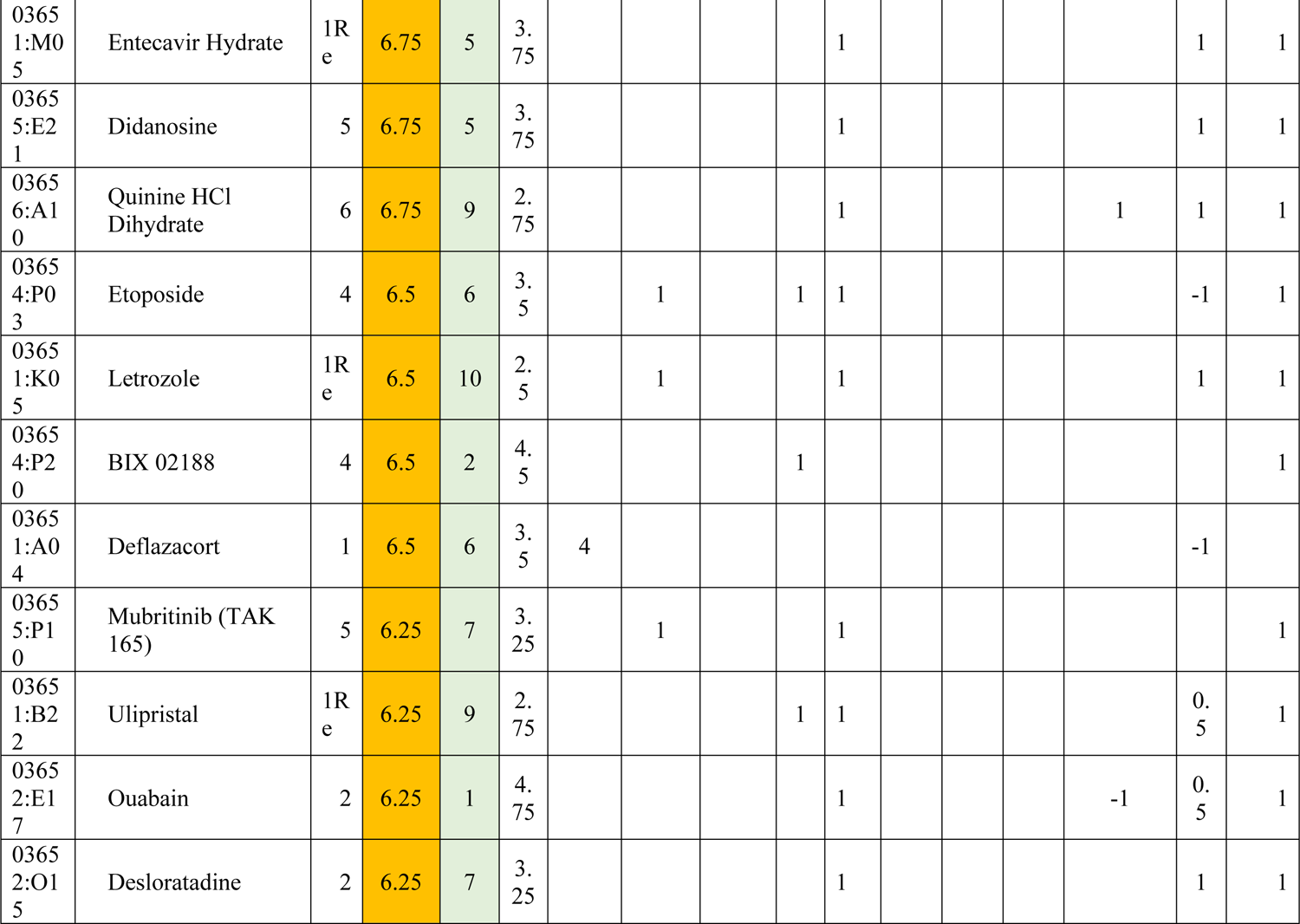
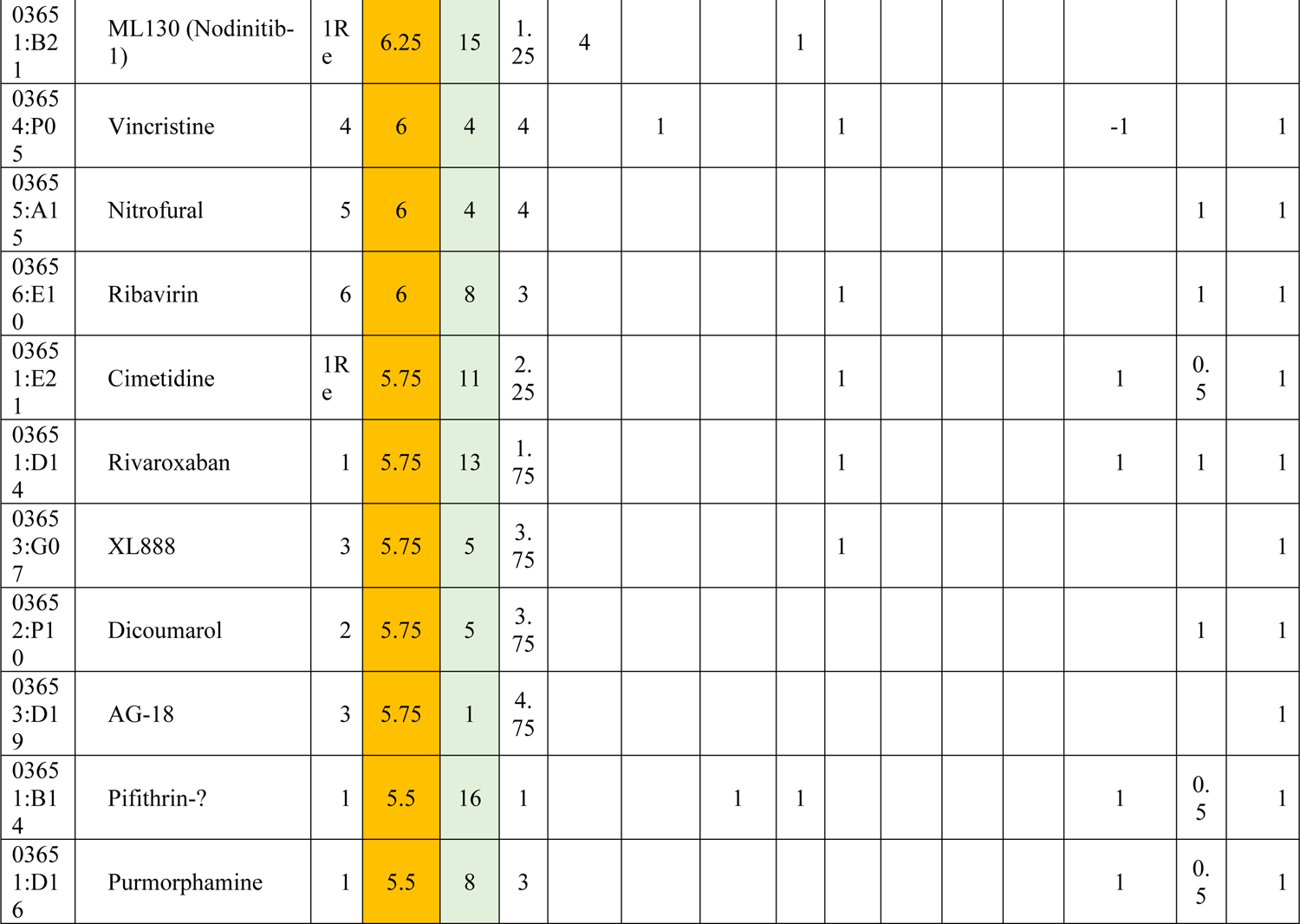
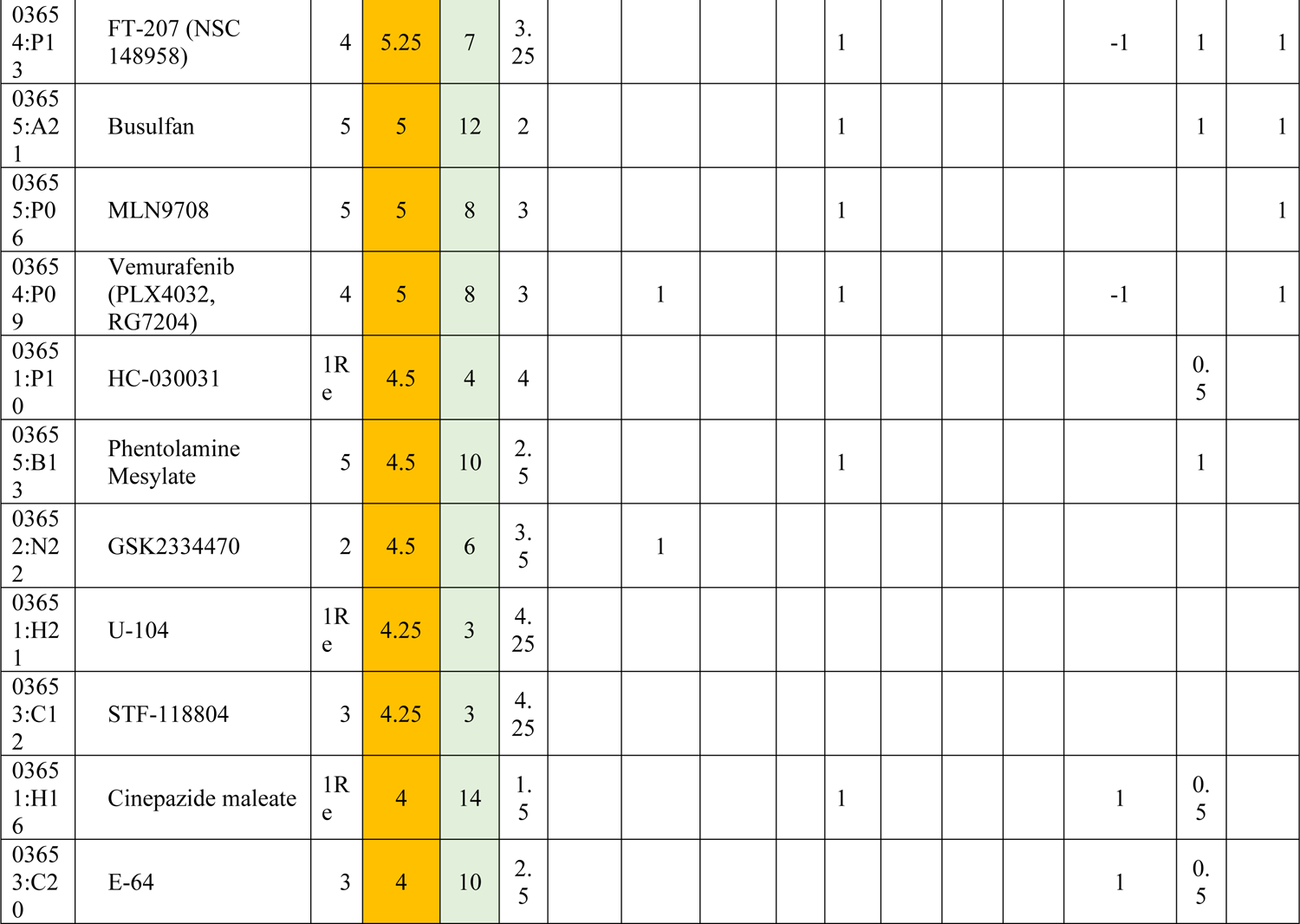
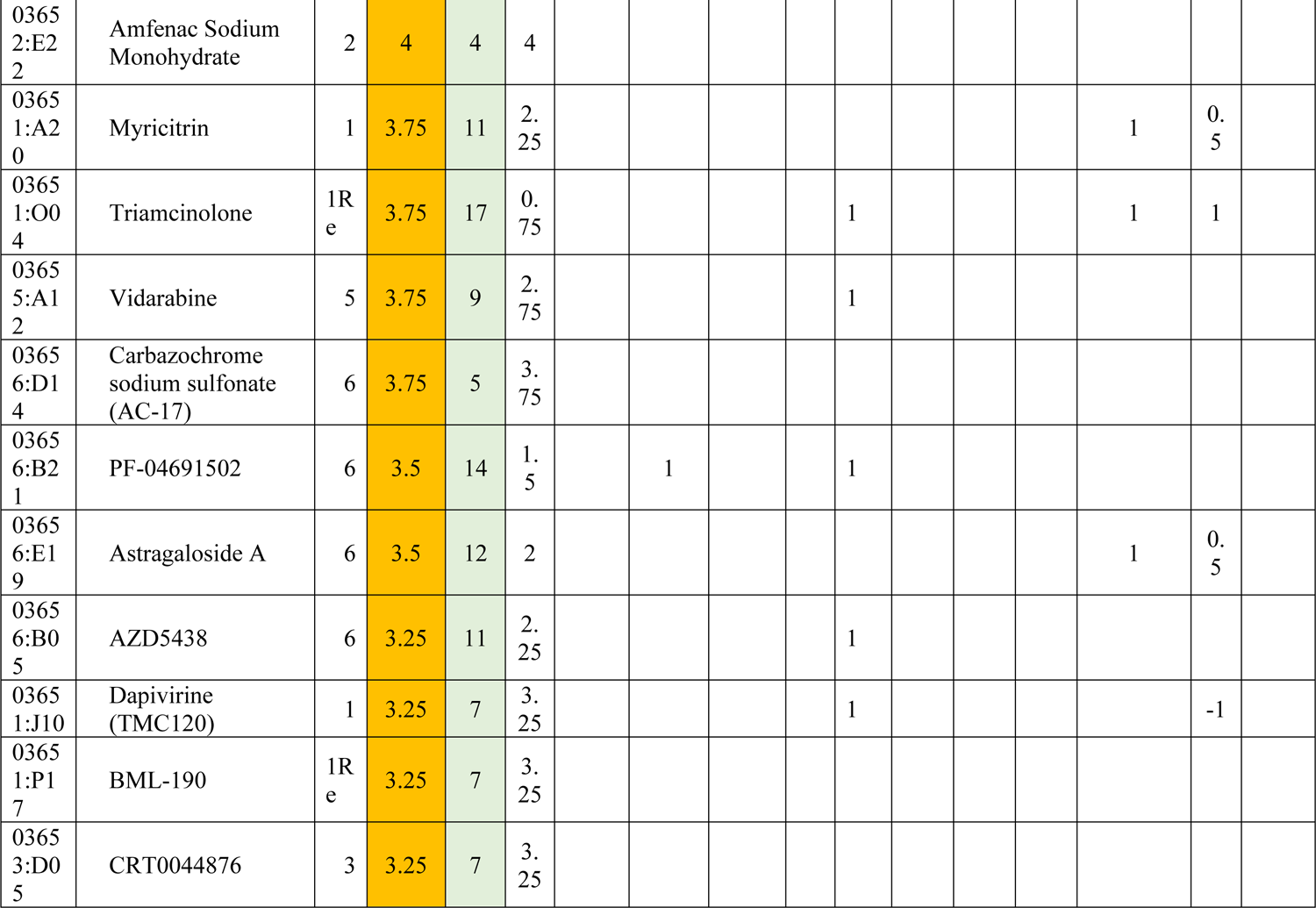
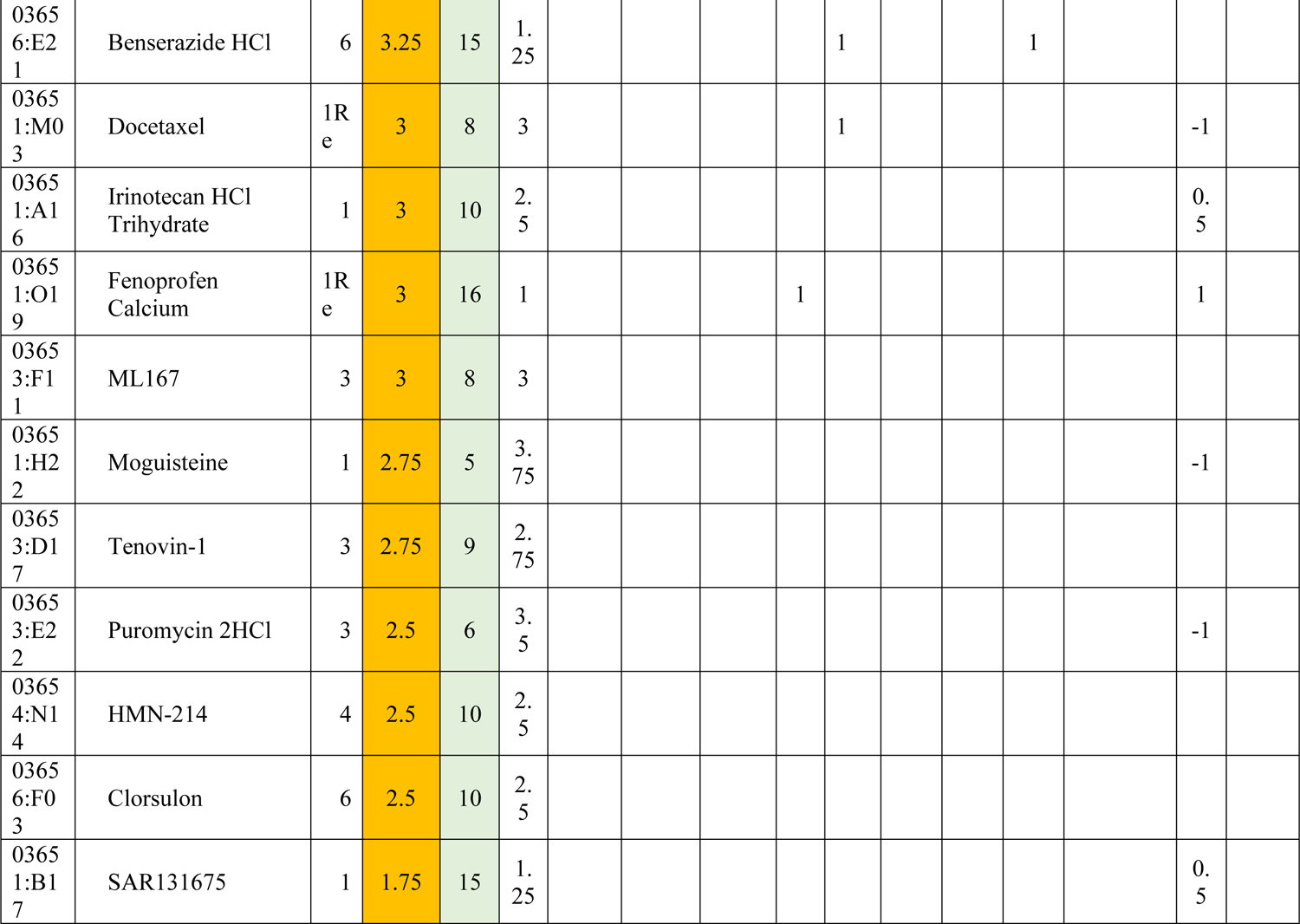
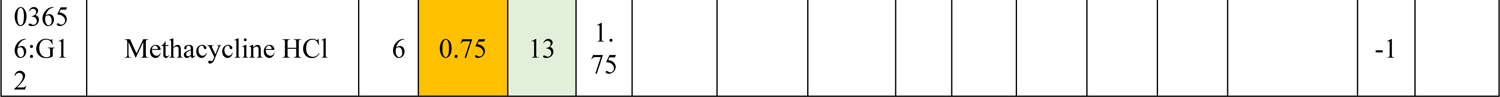
Selection of compounds for further validation.

**Supplemental Table 2:** Validation results for 44 selected compounds in primary rat neurons and human NGN2-iNs.

24h treatment, rat neurons
| Drug | Average fold change | P value (two tailed) | P value summary | Significant (alpha<0.05)? |
| --- | --- | --- | --- | --- |
| Forskolin | 3.639 | <0.0001 | **** | Yes |
| BIX 02188 | 3.175 | 0.011 | * | Yes |
| Ouabain | 3.06 | 0.0001 | *** | Yes |
| Digoxin | 2.707 | 0.0014 | ** | Yes |
| Nitazoxanide | 1.567 | 0.0012 | ** | Yes |
| Rigosertib (ON-01910) | 1.221 | 0.0173 | * | Yes |
| Pelitinib (EKB-569) | 1.199 | 0.0185 | * | Yes |
| Letrozole | 1.198 | 0.0302 | * | Yes |
| Rocilinostat (ACY-1215) | 1.163 | 0.0296 | * | Yes |
| Quercetin Dihydrate | 1.127 | 0.0492 | * | Yes |
| Betahistine 2HCl | 1.127 | 0.0134 | * | Yes |
| Amoxicillin Sodium | 1.105 | 0.039 | * | Yes |
| Edaravone | 1.081 | 0.0134 | * | Yes |
| Almotriptan | 0.9284 | 0.0344 | * | Yes |
| Etoposide | 1.206 | 0.1674 | ns | No |
| XL888 | 1.192 | 0.0649 | ns | No |
| Didanosine | 1.084 | 0.2885 | ns | No |
| U-104 | 1.078 | 0.5031 | ns | No |
| FT-207 (NSC 148958) | 1.069 | 0.5568 | ns | No |
| Budesonide | 1.065 | 0.5814 | ns | No |
| AG-18 | 1.064 | 0.1908 | ns | No |
| LY2811376 | 1.061 | 0.1923 | ns | No |
| Aminoglutethimide | 1.057 | 0.3665 | ns | No |
| Apixaban | 1.053 | 0.1083 | ns | No |
| Azaperone | 1.049 | 0.3419 | ns | No |
| Entecavir Hydrate | 1.046 | 0.3067 | ns | No |
| Inosine | 1.042 | 0.5278 | ns | No |
| Roflumilast | 1.03 | 0.4851 | ns | No |
| Ulipristal | 1.021 | 0.7969 | ns | No |
| Azelastine HCl | 1.02 | 0.6803 | ns | No |
| Desloratadine | 1.018 | 0.7573 | ns | No |
| PJ34 | 1.017 | 0.8487 | ns | No |
| Empagliflozin (BI 10773) | 1.004 | 0.9457 | ns | No |
| Deflazacort | 1.003 | 0.9635 | ns | No |
| Dabrafenib (GSK2118436) | 0.9891 | 0.3583 | ns | No |
| Nitrofurantoin | 0.9846 | 0.6657 | ns | No |
| Glycyrrhizic Acid | 0.9801 | 0.8447 | ns | No |
| Ribavirin | 0.9731 | 0.6684 | ns | No |
| Rolipram | 0.972 | 0.3145 | ns | No |
| Azilsartan | 0.9716 | 0.6438 | ns | No |
| Dicoumarol | 0.9577 | 0.5499 | ns | No |
| ML130 (Nodinitib-1) | 0.9557 | 0.4339 | ns | No |
| HC-030031 | 0.9248 | 0.265 | ns | No |
| URB597 | 0.9235 | 0.5509 | ns | No |
| Mubritinib (TAK 165) | 0.9158 | 0.1805 | ns | No |
| DMSO | 1 | N/A | N/A | N/A |

72h treatment, rat neurons
| Drug | Average fold change | P value (two tailed) | P value summary | Significant (alpha<0.05)? |
| --- | --- | --- | --- | --- |
| Forskolin | 5.172 | <0.0001 | **** | Yes |
| BIX 02188 | 3.787 | 0.0002 | *** | Yes |
| Ouabain | 2.674 | 0.0015 | ** | Yes |
| Digoxin | 2.204 | 0.0033 | ** | Yes |
| Pelitinib (EKB-569) | 1.838 | 0.0097 | ** | Yes |
| XL888 | 1.715 | 0.0335 | * | Yes |
| Etoposide | 1.552 | 0.0077 | ** | Yes |
| Rocilinostat (ACY-1215) | 1.411 | 0.0049 | ** | Yes |
| Nitazoxanide | 1.386 | 0.0222 | * | Yes |
| Budesonide | 1.287 | 0.0014 | ** | Yes |
| LY2811376 | 1.129 | 0.0178 | * | Yes |
| Aminogluthethimide | 0.8203 | 0.0374 | * | Yes |
| Rigosertib (ON-01910) | 1.357 | 0.0652 | ns | No |
| Letrozole | 1.192 | 0.0645 | ns | No |
| Quercetin Dihydrate | 1.159 | 0.0635 | ns | No |
| Mubritinib (TAK 165) | 1.147 | 0.2364 | ns | No |
| Empagliflozin (BI 10773) | 1.141 | 0.0916 | ns | No |
| Desloratadine | 1.132 | 0.1516 | ns | No |
| Azelastine HCl | 1.113 | 0.1408 | ns | No |
| Ulipristal | 1.112 | 0.1648 | ns | No |
| Rolipram | 1.088 | 0.2542 | ns | No |
| Apixaban | 1.087 | 0.0924 | ns | No |
| U-104 | 1.079 | 0.3396 | ns | No |
| Betahistine 2HCl | 1.073 | 0.2481 | ns | No |
| Edaravone | 1.061 | 0.2385 | ns | No |
| Entecavir Hydrate | 1.055 | 0.3392 | ns | No |
| Glycyrrhizic Acid | 1.037 | 0.6864 | ns | No |
| Amoxicillin Sodium | 1.032 | 0.7329 | ns | No |
| Nitrofurantoin | 1.029 | 0.8807 | ns | No |
| Deflazacort | 1.018 | 0.8445 | ns | No |
| Didanosine | 1.017 | 0.8106 | ns | No |
| FT-207 (NSC 148958) | 1.017 | 0.6764 | ns | No |
| Ribavirin | 1.007 | 0.9078 | ns | No |
| Azaperone | 0.9961 | 0.9079 | ns | No |
| Dicoumarol | 0.9762 | 0.7672 | ns | No |
| AG-18 | 0.9666 | 0.5969 | ns | No |
| Azilsartan | 0.9656 | 0.6346 | ns | No |
| Dabrafenib (GSK2118436) | 0.9353 | 0.2136 | ns | No |
| Inosine | 0.9109 | 0.1424 | ns | No |
| URB597 | 0.9092 | 0.1946 | ns | No |
| Almotriptan | 0.8949 | 0.0799 | ns | No |
| HC-030031 | 0.8796 | 0.1298 | ns | No |
| ML130 (Nodinitib-1) | 0.8795 | 0.1024 | ns | No |
| Roflumilast | 0.8752 | 0.1591 | ns | No |
| PJ34 | 0.8582 | 0.0887 | ns | No |
| DMSO | 1 | N/A | N/A | N/A |

24h treatment, human NGN2-iNs
| Drug | Average fold change | P value (two tailed) | P value summary | Significant (alpha<0.05)? |
| --- | --- | --- | --- | --- |
| Digoxin | 2.86 | 0.0251 | * | Yes |
| Ouabain | 2.425 | 0.0129 | * | Yes |
| Rigosertib (ON-01910) | 1.663 | 0.0206 | * | Yes |
| Forskolin | 1.391 | 0.0153 | * | Yes |
| Budesonide | 1.199 | 0.0355 | * | Yes |
| Didanosine | 1.578 | 0.1375 | ns | No |
| Empagliflozin (BI 10773) | 1.534 | 0.2076 | ns | No |
| Quercetin Dihydrate | 1.437 | 0.3354 | ns | No |
| Azelastine HCl | 1.378 | 0.1517 | ns | No |
| Dabrafenib (GSK2118436) | 1.378 | 0.381 | ns | No |
| Deflazacort | 1.338 | 0.2725 | ns | No |
| Betahistine 2HCl | 1.326 | 0.2401 | ns | No |
| Nitazoxanide | 1.324 | 0.1063 | ns | No |
| Almotriptan | 1.311 | 0.2891 | ns | No |
| AG-18 | 1.266 | 0.0557 | ns | No |
| Glycyrrhizic Acid | 1.265 | 0.461 | ns | No |
| Inosine | 1.234 | 0.2321 | ns | No |
| BIX 02188 | 1.228 | 0.2411 | ns | No |
| Entecavir Hydrate | 1.221 | 0.2515 | ns | No |
| Letrozole | 1.217 | 0.2271 | ns | No |
| Rolipram | 1.182 | 0.224 | ns | No |
| Aminoglutethimide | 1.177 | 0.1727 | ns | No |
| Etoposide | 1.173 | 0.3406 | ns | No |
| URB597 | 1.168 | 0.1867 | ns | No |
| U-104 | 1.163 | 0.4164 | ns | No |
| Nitrofurantoin | 1.145 | 0.3707 | ns | No |
| ML130 (Nodinitib-1) | 1.143 | 0.2225 | ns | No |
| FT-207 (NSC 148958) | 1.133 | 0.4391 | ns | No |
| Rocilinostat (ACY-1215) | 1.111 | 0.4474 | ns | No |
| Roflumilast | 1.104 | 0.6772 | ns | No |
| Amoxicillin Sodium | 1.102 | 0.1956 | ns | No |
| XL888 | 1.099 | 0.4062 | ns | No |
| Azilsartan | 1.093 | 0.5713 | ns | No |
| Pelitinib (EKB-569) | 1.089 | 0.5371 | ns | No |
| Azaperone | 1.079 | 0.4025 | ns | No |
| PJ34 | 1.056 | 0.6615 | ns | No |
| Edaravone | 1.043 | 0.5514 | ns | No |
| Dicoumarol | 1.041 | 0.8104 | ns | No |
| Ulipristal | 1.039 | 0.7041 | ns | No |
| Apixaban | 1.023 | 0.764 | ns | No |
| Desloratadine | 1.023 | 0.8387 | ns | No |
| HC-030031 | 1.023 | 0.8394 | ns | No |
| LY2811376 | 1.007 | 0.9426 | ns | No |
| Ribavirin | 0.9954 | 0.9648 | ns | No |
| Mubritinib (TAK 165) | 0.9671 | 0.7106 | ns | No |
| DMSO | 1 | N/A | N/A | N/A |

**Supplemental Table 3:**
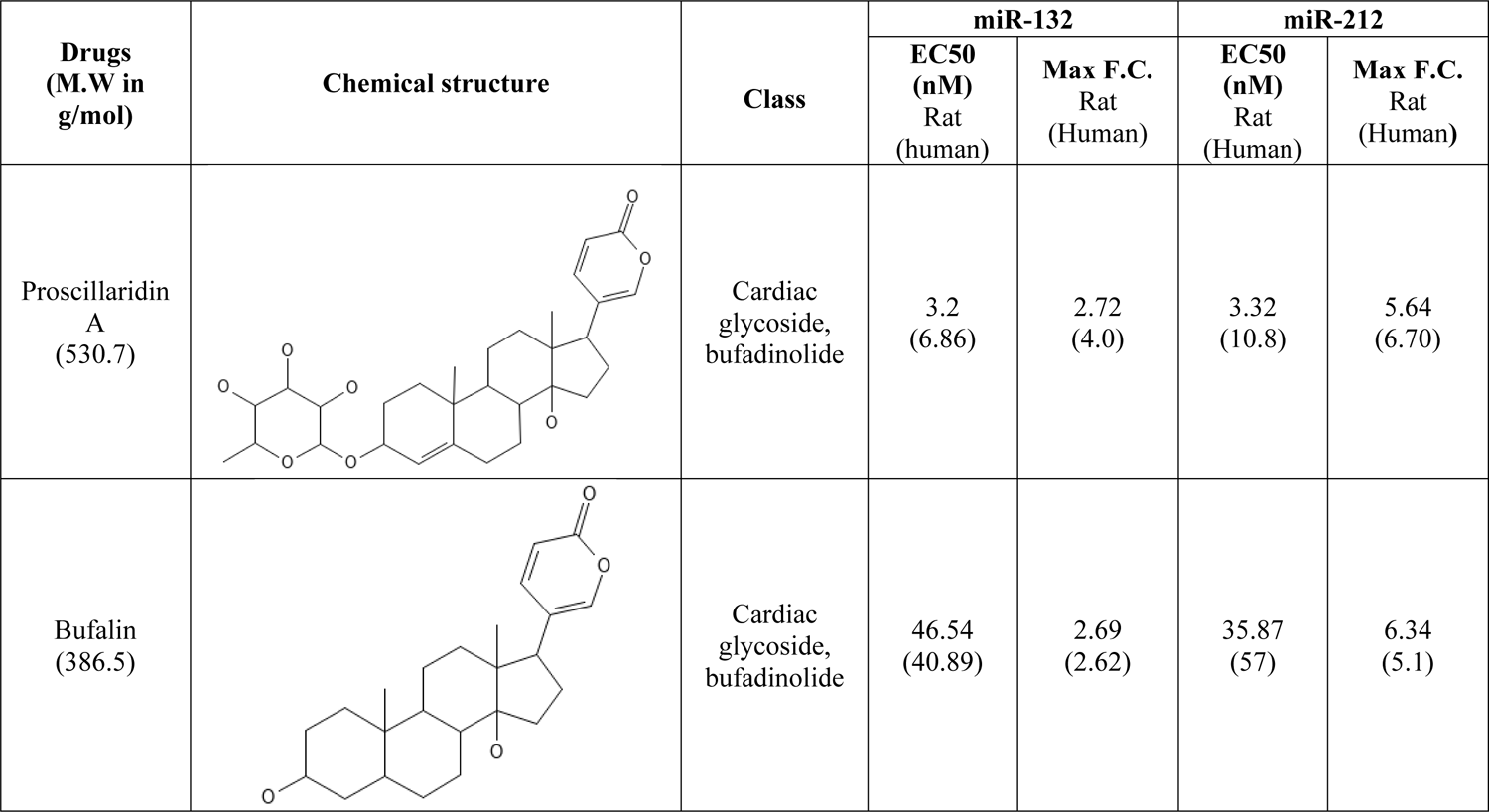
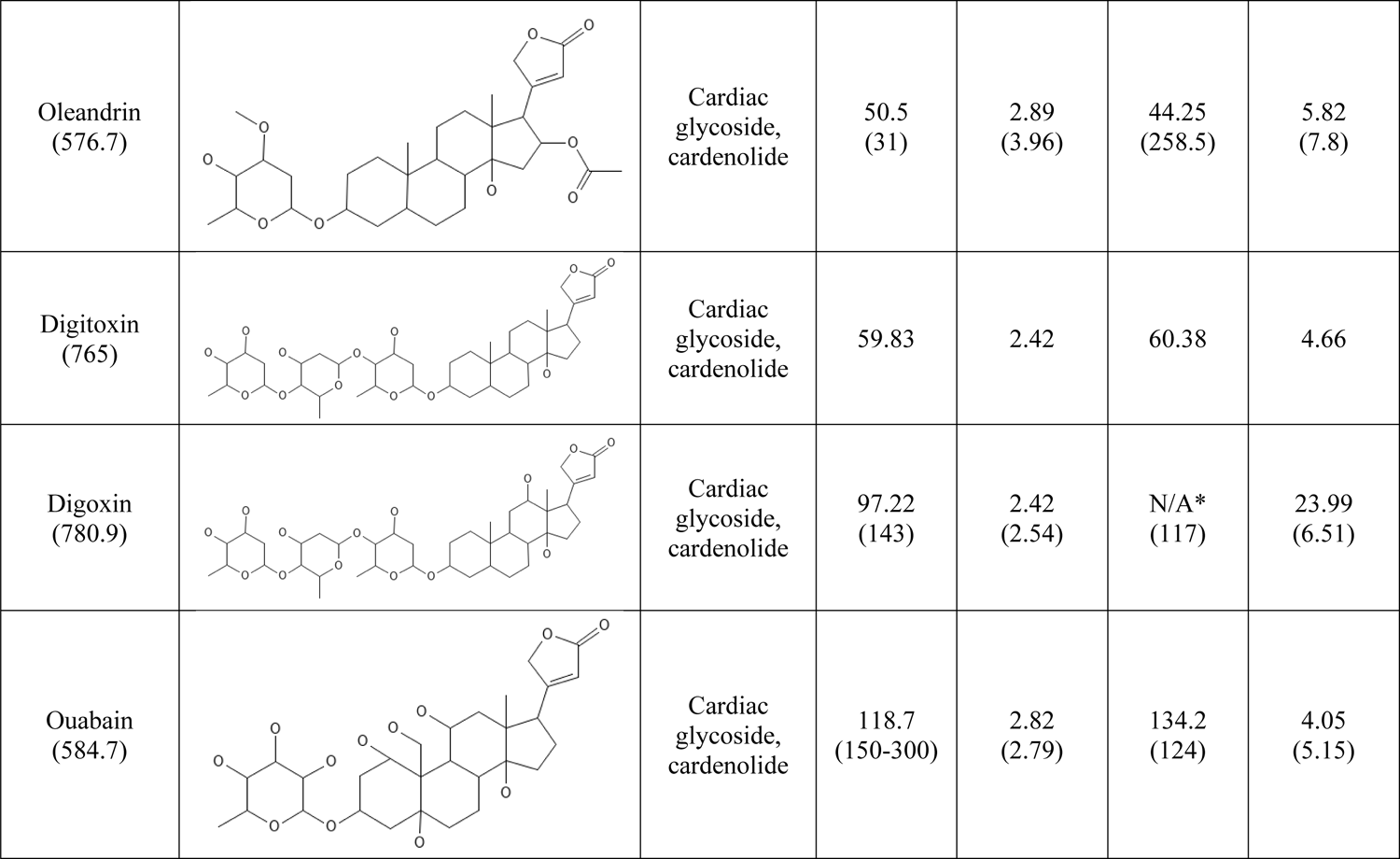
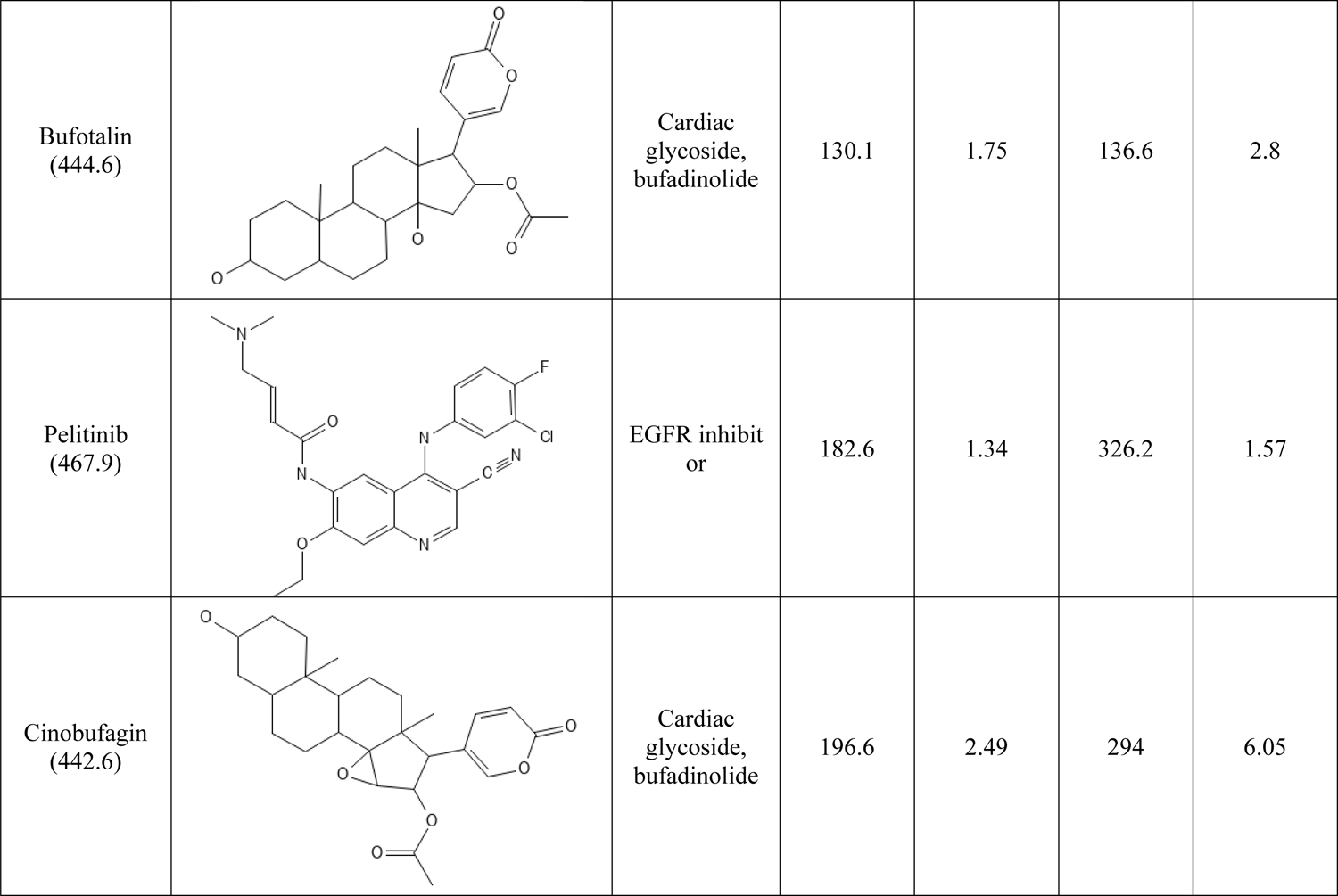
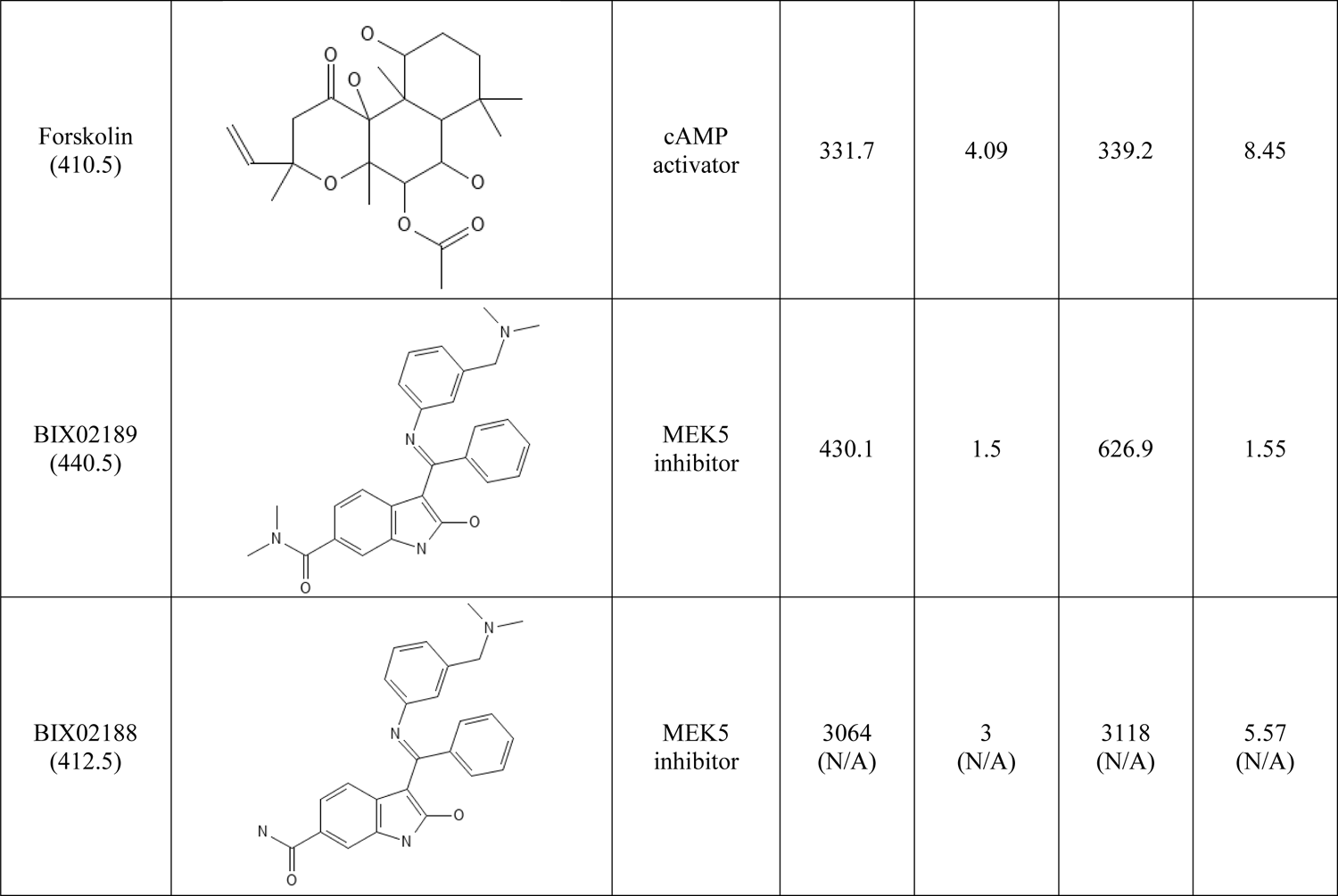
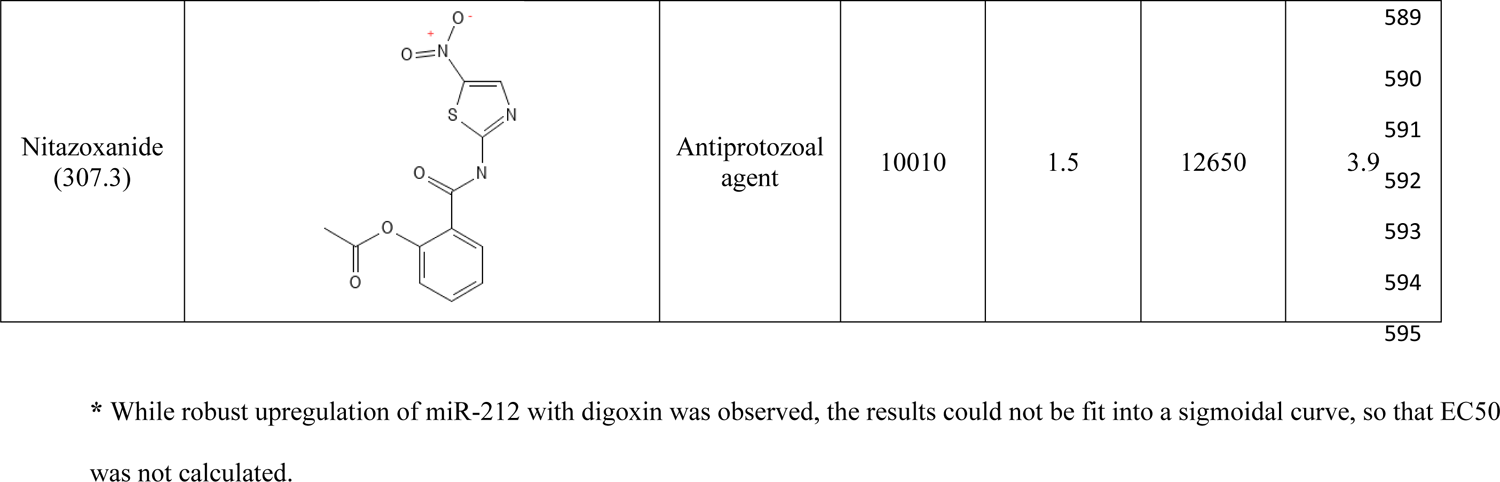
Chemical structures and miR-132/212 EC_50_ and max fold change of various compounds.

**Supplemental Table 4:**
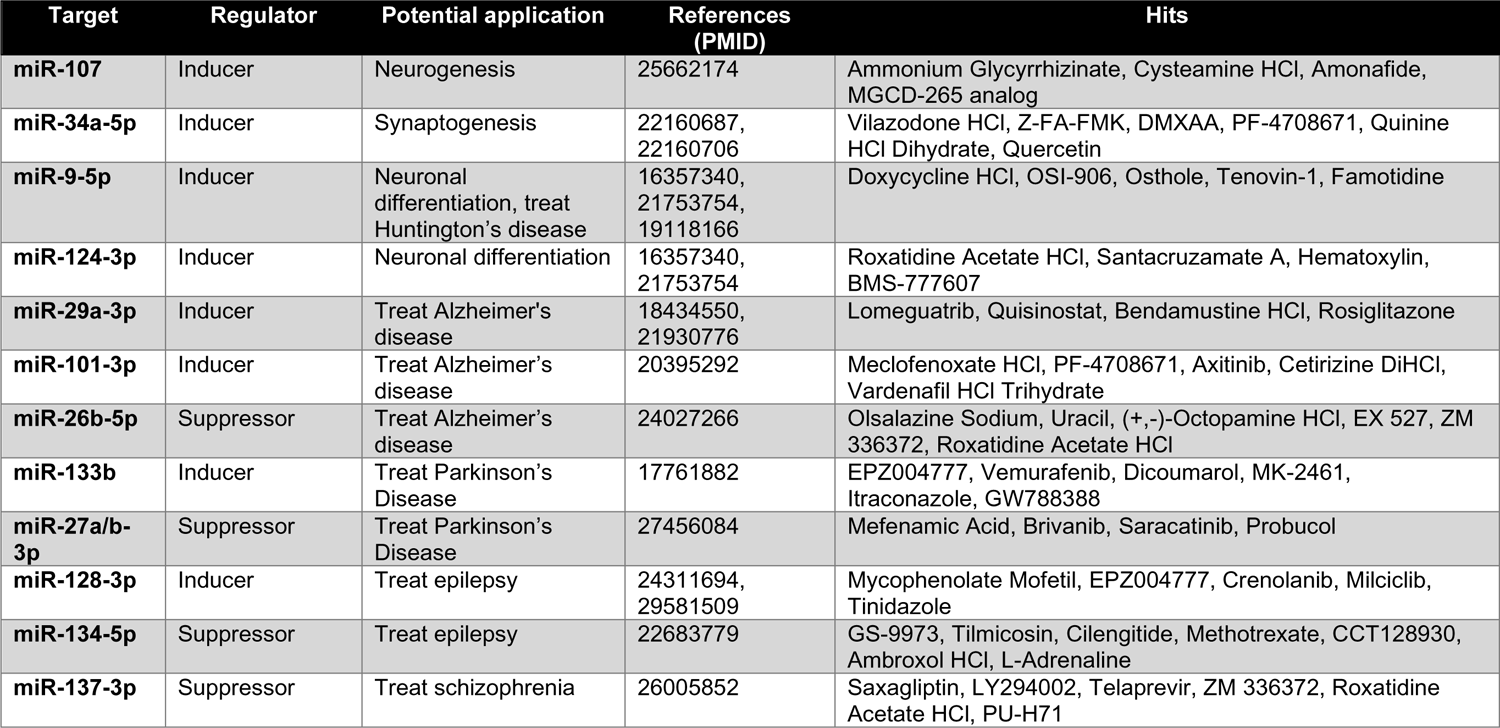
Candidate small molecule compounds that may regulate miRNAs implicated in neurological disease.

**Supplemental Table 5:** List of commercial miRNA primers used for miRNA RT-qPCR.

| No. | Name | GeneGlobe ID | Species |
| --- | --- | --- | --- |
| 1 | hsa-let-7a-5p | YP00205727 | Human, mouse, rat |
| 2 | hsa-mir-22-3p | YP00204606 | Human, mouse, rat |
| 3 | hsa-miR-26b-5p | YP00204172 | Human, mouse, rat |
| 4 | hsa-miR-99a-5p | YP00204521 | Human, mouse, rat |
| 5 | hsa-miR-103a-3p | YP00204063 | Human, mouse, rat |
| 6 | mmu-miR-124-3p | YP02119832 | Mouse, rat |
| 7 | hsa-miR-125a-5p | YP00204339 | Human, mouse, rat |
| 8 | hsa-miR-128-3p | YP00205995 | Human, mouse, rat |
| 9 | hsa-miR-132-3p | YP00206035 | Human, mouse, rat |
| 10 | hsa-miR-138-1-3p | YP00205881 | Human, mouse, rat |
| 11 | hsa-miR-191-5p | YP00204306 | Human, mouse, rat |
| 12 | hsa-miR-212-3p | YP00204170 | Human |
| 13 | mmu-miR-212-3p | YP00206022 | Mouse, rat |
| 14 | hsa-miR-107 | YP00204468 | Human, mouse, rat |

**Supplemental Table 6:**
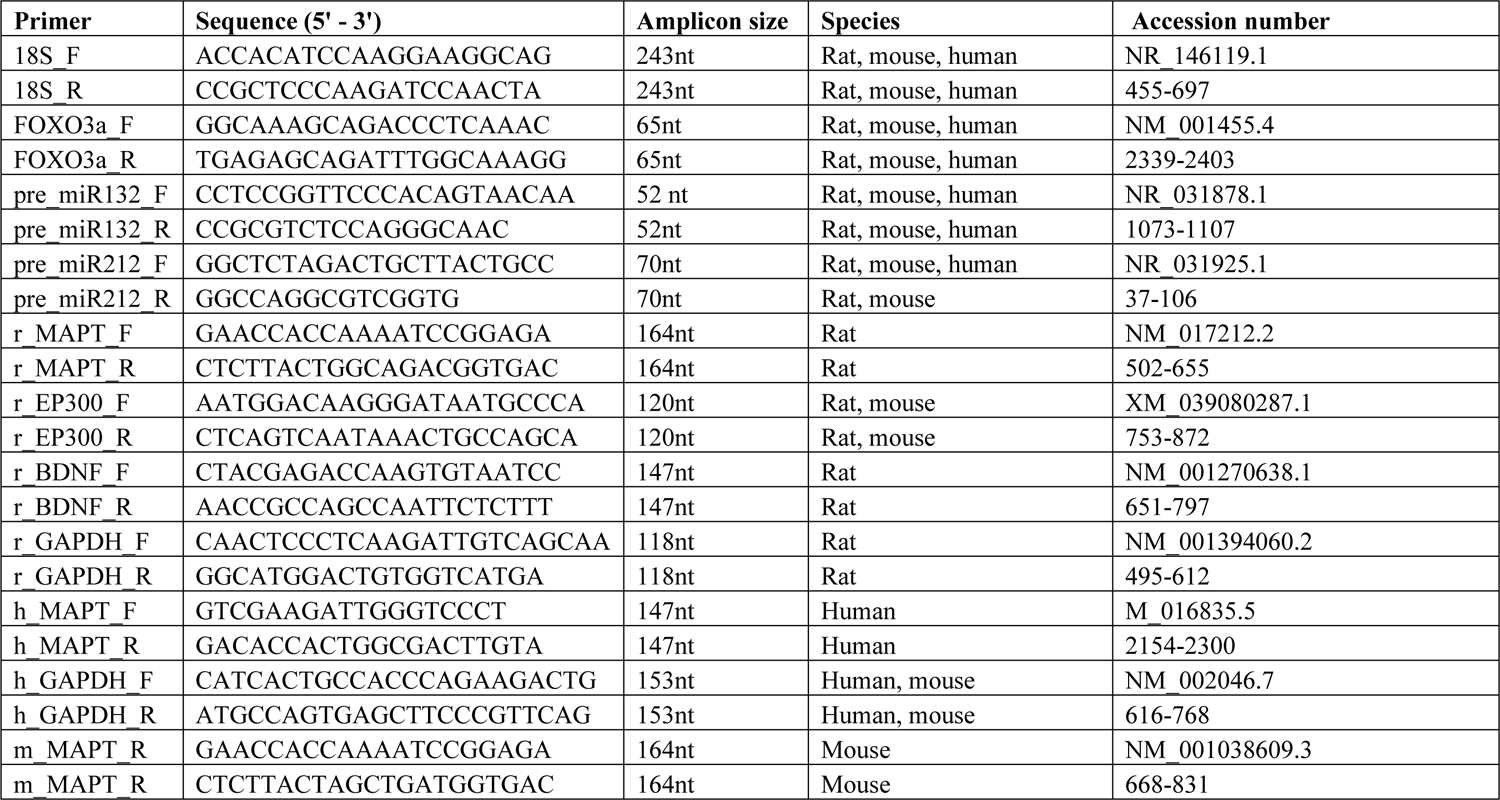
List of mRNA primer sequences used for mRNA RT-qPCR.

**Supplemental Table 7:**
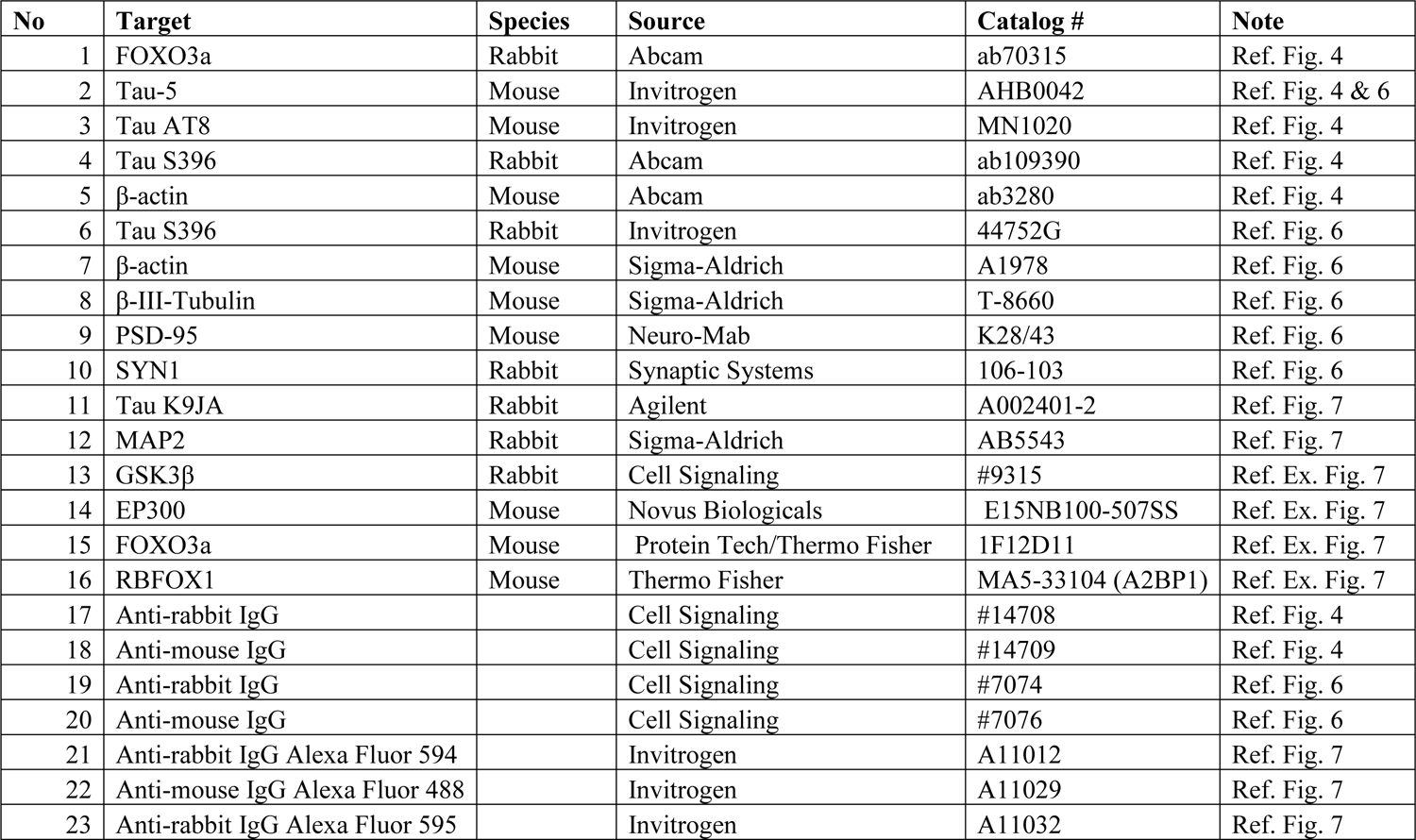
List of primary and secondary antibodies used for Western blots.

| No | Target | Species | Source | Catalog # | Note |
| --- | --- | --- | --- | --- | --- |
| 1 | FOXO3a | Rabbit | Abcam | ab70315 | Ref. Fig. 4 |
| 2 | Tau-5 | Mouse | Invitrogen | AHB0042 | Ref. Fig. 4 & 6 |
| 3 | Tau AT8 | Mouse | Invitrogen | MN1020 | Ref. Fig. 4 |
| 4 | Tau S396 | Rabbit | Abcam | ab109390 | Ref. Fig. 4 |
| 5 | β-actin | Mouse | Abcam | ab3280 | Ref. Fig. 4 |
| 6 | Tau S396 | Rabbit | Invitrogen | 44752G | Ref. Fig. 6 |
| 7 | β-actin | Mouse | Sigma-Aldrich | A1978 | Ref. Fig. 6 |
| 8 | β-III-Tubulin | Mouse | Sigma-Aldrich | T-8660 | Ref. Fig. 6 |
| 9 | PSD-95 | Mouse | Neuro-Mab | K28/43 | Ref. Fig. 6 |
| 10 | SYN1 | Rabbit | Synaptic Systems | 106-103 | Ref. Fig. 6 |
| 11 | Tau K9JA | Rabbit | Agilent | A002401-2 | Ref. Fig. 7 |
| 12 | MAP2 | Rabbit | Sigma-Aldrich | AB5543 | Ref. Fig. 7 |
| 13 | GSK3β | Rabbit | Cell Signaling | #9315 | Ref. Ex. Fig. 7 |
| 14 | EP300 | Mouse | Novus Biologicals | E15NB100-507SS | Ref. Ex. Fig. 7 |
| 15 | FOXO3a | Mouse | Protein Tech/Thermo Fisher | 1F12D11 | Ref. Ex. Fig. 7 |
| 16 | RBFOX1 | Mouse | Thermo Fisher | MA5-33104 (A2BP1) | Ref. Ex. Fig. 7 |
| 17 | Anti-rabbit IgG |  | Cell Signaling | #14708 | Ref. Fig. 4 |
| 18 | Anti-mouse IgG |  | Cell Signaling | #14709 | Ref. Fig. 4 |
| 19 | Anti-rabbit IgG |  | Cell Signaling | #7074 | Ref. Fig. 6 |
| 20 | Anti-mouse IgG |  | Cell Signaling | #7076 | Ref. Fig. 6 |
| 21 | Anti-rabbit IgG Alexa Fluor 594 |  | Invitrogen | A11012 | Ref. Fig. 7 |
| 22 | Anti-mouse IgG Alexa Fluor 488 |  | Invitrogen | A11029 | Ref. Fig. 7 |
| 23 | Anti-rabbit IgG Alexa Fluor 595 |  | Invitrogen | A11032 | Ref. Fig. 7 |

## MATERIALS AND METHODS

### Induced neuron differentiation from iPSC

Induced pluripotent stem cell (iPSC) lines were retrieved and differentiated into neurons with NGN2 expression, as previously reported^23^. Briefly, iPSCs were plated in mTeSR1 media at a density of 9.5×10^4^ cells/cm2 on Matrigel (Corning #354234)-coated plates. Cells were then transduced with the following virus: pTet-O-NGN2-puro (Addgene #52047): 0.1 μL per 5×10^4^ cells; Tet-O-FUW-eGFP (Addgene #30130): 0.05μL per 5×10^4^ cells; Fudelta GW-rtTA (Addgene #19780): 0.11 μL per 5×10^4^ cells. Transduced cells were dissociated with Accutase (StemCell Technologies) and plated onto Matrigel-coated plates in mTeSR1 (StemCell Technologies) at 5×10^4^/cm^2^ (Day 0). On day 1, media was changed to KSR media (Knockout DMEM, 15% KOSR, 1x MEM-NEAA, 55 μM beta-mercaptoethanol, 1x GlutaMAX; Gibco) with doxycycline (2 μg/ml, Sigma-Aldrich). Doxycyline was maintained in the media for the remainder of the differentiation. On day 2, the media was changed to 1:1 KSR: N2B media with puromycin (5 μg/ml, Gibco), where N2B was composed of DMEM/F12, 1x GlutaMAX, 1x N2 supplement B (StemCell Technologies) and 0.3% dextrose (Sigma-Aldrich). Puromycin was maintained in the media throughout the differentiation. On day 3, the media was changed to N2B media + 1:100 B-27 supplement (GIBCO) and puromycin (10 μg/ml). From day 4 on, cells were cultured in NBM media (Neurobasal medium, 0.5x MEM-NEAA, 1x GlutaMAX, 0.3% dextrose) + 1:50 B-27 + BDNF, GDNF, CNTF (10 ng/ml each, Peprotech). After day 4, half of the media was replaced by fresh media twice per week. Cells were stocked on day 4 at 1∼2×10^6^ cells in 200 μL freezing media (50% day 4 media + 40% FBS + 10% DMSO) per cryovial in −80°C overnight, followed by liquid nitrogen storage. iPSC-derived neurons used for validation experiments were prepared similarly. iPSC lines were generated following review and approval through Brigham and Women’s Hospital Institutional Review Board (IBR#2015P001676).

### Preparation of NGN2-iNs for high-throughput screen

Twenty-five 96-well plates (Corning) were coated with Matrigel solution (0.2 mg/mL in DMEM/F12) at 60 μL per well for 1.5 hours at 37°C. Then, the Matrigel solution was completely removed, and 100 μL PBS (Gibco) was added per well using electronic 12-channel pipettes in speed 3 (e12c-pip; Eppendorf). The plates were temporality incubated at 37°C. Frozen day 4 iPSC-iNs were thawed in 500 μL pre-warmed resuspension media per vial, which was composed of NBM, 1:100 B27, and 1:1000 ROCKi (StemCell Technologies), and were kept in a warm metal bath to facilitate the thawing. Multiple vials were pooled into one 50 mL conical tube, then pre-warmed resuspension media were added drop-wisely to reach the volume of 40 mL. After gently mixing by reverting the tube, viable cell concentration was counted with trypan blue (Bio-Rad). The cells were spun down (220 g, 5 min, room temperature) and resuspended in day 4 media at 1×10^5^ cells/mL. Then, PBS was completely removed from the plates, and 100 μL cell suspension was added per well, using e12c-pip at speed 3. The cells were incubated at 37°C after shaking the plates for even distribution (day 4). To reduce evaporation during incubation, plates were kept in plastic containers lined with sterile wet paper towels. On day 5, an additional 100 μL pre-warmed day 4 medium was added per well using e12c-pip in speed 1. On days 7/10/14/18, 95 μL conditioned medium was removed, and 100 μL pre-warmed day 4 medium was added per well, both using e12c-pip.

### High-throughput screen in NGN2-iNs

On day 19, half (three 384-well plates) of the Selleck bioactive compound library (N=1902 compounds) were pin transferred (V&P Scientific) to twelve NGN2-iN plates using the Seiko Compound Transfer Robot at 200 nL per well (final concentration at 10 μM). Positive control (Forskolin) and negative control (DMSO) were also pin transferred to the wells without library compound. On day 20, four 10x photos were taken per well automatically using the ImageXpress Micro Confocal microscope (Molecular Devices). Then, the media in the plates was removed with approximately 20 μL media left, using a 24-channel stainless steel manifold (Drummond #3-000-101) linked with a vacuum at a low speed. With the help of Multidrop™ Combi Reagent Dispenser (Thermo Scientific) and the standard cassette (speed: low), 250 μL ice-cold DPBS (Wisent) was added per well. The DPBS was removed with approximately 20 μL liquid left, using another 24-channel stainless steel manifold linked with the vacuum at a low speed. The residual DPBS was completely removed using a mechanic 12-channel pipette. Next, 45 μL lysing solution was added per well using another Multidrop™ Combi Reagent Dispenser with the small cassette (speed: low), where the lysing solution is composed of single-cell lysis buffer (Takara #635013): 1x RNase Inhibitor Murine (NEB): nuclease-free water (Exiqon) = 19:1:190. Thorough lysis was achieved by shaking the plates on a shaker for 5 min at room temperature. The lysis samples were transferred from four 96-well plates to each 384-square-well plate using the 96-well module-coupled VPrep liquid handler (Agilent) with 30 μL tips (twice without changing tips). After sealing, the 384-square-well plates were spun down at 4000 rpm for 5 min, and 10 μL supernatant was aliquoted to a 384-well plate (Eppendorf) using the 384-well module-coupled VPrep liquid handler. The plates were finally sealed with the PlateLoc Heat Sealer (Agilent) and stored at −20°C. The other half (three and a quarter 384-well plates) of the compound library were added to the remaining thirteen 96-well plates after one day delay (on day 20), using the identical protocol due to the time consumption. The high-throughput screen was conducted in the ICCB-Longwood Screening Facility, Harvard Medical School.

### Ultra-low input miRNA-seq using the RealSeq

To avoid RNA purification, we used RealSeq-T technology (RealSeq Biosciences) following manufacturer recommendations. In summary, cell lysates were incubated at 70C for 5 minutes on RealSeq hybridization buffer (100 mM NaCl, 50 mM Tris-HCl, 10 mM MgCl2, 1mM DTT, pH 7.9) with 1x RealSeq biotinylated DNA probes to target all miRNAs in miRbase 21. After 2 hours of incubation at 37C, 10 μL of RealSeq Beads were added, and miRNAs were captured using a 384-well Magnet Plate (Alpaqua, MA). Following three washes with RealSeq Wash buffer, miRNA was eluted from beads in 10uL of RNase-free water. All the miRNA elusion was input to prepare sequencing libraries with RealSeq-Biofluids following manufacturer instructions (RealSeq Biosciences). In summary, a single adapter and circularization approach was used ^24^. Libraries were barcoded with dual indexes and sequenced with a NextSeq 550 (Illumina, CA). FastQ files were trimmed of adapter sequences using Cutadapt with the following parameters: cutadapt -u 1 - a TGGAATTCTCGGGTGCCAAGG-m 15. Trimmed reads were aligned to the corresponding reference by using Bowtie ^59^. Counts of each miRNA were normalized among samples by total miRNA read counts.

### Animal Use

This study was carried out in accordance with the recommendations in the U.S. National Institutes of Health Guide for the Care and Use of Laboratory Animals. The protocol was approved by the Institutional Animal Care and Use Committee at Brigham and Women’s Hospital. Mice were maintained on a 12:12-h light/dark cycle (7:00 am on/7:00 pm off) with food and water provided ad libitum before experimental procedures

### Rat and Mouse Primary Neuron Culture

Rat primary cortical neuron cultures were prepared from E18 SAS Sprague Dawley pups (Charles River). Brain tissues were dissected, dissociated enzymatically by 0.25% Trypsin-EDTA (Thermo Fisher Scientific), triturated with fire-polished glass Pasteur pipettes, and passed through a 40 μm cell strainer (Sigma-Aldrich) to remove clumps. After counting, neurons were seeded onto poly-D-lysine (Sigma-Aldrich) coated cell culture plates at 80,000 cells/cm^2^ in neurobasal medium supplemented with 1X B27 and 0.25X GlutaMax. Half of the cell medium was changed every 4 days until use.

Mouse primary cortical neuron cultures were prepared from P1 or P2 postnatal pups from PS19 mouse breeding pairs. After dissection, mouse brain tissues were kept in Hibernate-A medium at 4°C in the dark for ∼4h. After genotyping, brains from pups of the same genotype, either WT or PS19, were pooled together and dissociated enzymatically with papain solution (Worthington). After dissociation, mouse neurons were prepared and cultured similarly to rat neurons.

### siRNA and miRNA Mimics Transfection

siRNAs and miRNA mimics were purchased from Dharmacon (Horizon Discovery) and were dissolved in nuclease-free water to prepare 50 μM stock concentrations. Transfection was performed using NeuroMag (OZ Biosciences). For siRNA knockdown, transfection was performed with 50 nM siRNAs on DIV7 and DIV9, and RNA was collected for analysis at DIV11. Transfection of DIV14 neurons with 50 nM miR-132 or CTRL mimics was performed similarly. RNA was collected for analysis 72h later at DIV17.

### RNA Extraction, cDNA Preparation, and RT-qPCR

Total RNA from cells was extracted using the Norgen Total RNA Purification Kit (Norgen Biotek) following the manufacturer’s protocol. DNAse1 was applied during RNA extraction to remove genomic DNA. RNA was eluted in nuclease-free water, and the concentration was measured using Nanodrop (Thermo Fisher Scientific).

For miRNA analysis, 50ng of RNA was reverse transcribed into cDNA using the miRCURY LNA RT kit (Qiagen). RT-qPCR mix was prepared using the miRCURY LNA SYBR Green PCR kit (Qiagen). qPCR was performed using the QuantStudio 7 Flex System. The cycling conditions were 95°C for 10 min, 50 cycles of 95°C for 15 s, and 60°C for 1 min following dissociation analysis. miRNA expression was normalized to the geometric mean of miR-103a and let7a unless stated otherwise in figure legends. For mRNA analysis, 250-1000 ng of RNA was reverse transcribed into cDNA using the High Capacity cDNA Reverse Transcription Kit (Thermo Fisher Scientific). RT-qPCR mix was prepared using the PowerUp SYBR Green Master Mix (Thermo Fisher Scientific). qPCR was performed using the QuantStudio 7 Flex System. The cycling conditions were 95°C for 10 min, 50 cycles of 95°C for 15 s, and 60°C for 1 min following dissociation analysis. mRNA expression was normalized to the geometric mean of 18S and GAPDH. Quantification was performed using the delta-delta Ct method. miRNA and mRNA primers used were listed in Supplemental Tables 5 and 6.

### Transcriptome profile by RNA-seq

After quality control by Agilent 2100 Bioanalyzer, the total RNA was used as input for library preparation by Novogene Co., Ltd, followed by high-throughput sequencing on Illumina HiSeq X with PE150 mode to produce approximately 20 M reads per sample. The reads were quality controlled with FastQC, trimmed with Trimmomatic, aligned with HiSat2 to hg38, and quantified with HTSeq-count using the Galaxy platform. Read counts were processed for differential expression analysis using the R package DEBrowser with DESeq2. Pathway analysis was performed by Enrichr. Promoter binding sites were extracted from JASPAR 2022 TFBS via the UCSC genome browser.

### Western Blot Analysis

Total protein was extracted using RIPA buffer (Boston Bioproducts) supplemented with Complete, Mini, EDTA-free Protease Inhibitor Cocktail (Millipore Sigma). Protein concentrations were determined using the Micro BCA Protein Assay Kit (Thermo Fisher Scientific). Equal amounts of protein were loaded, and electrophoresis was performed in NuPAGE 4 to 12% gradient Bis-Tris polyacrylamide protein gels (Thermo Fisher Scientific). Proteins were transferred to Immun-Blot PVDF membranes (Bio-Rad) and then blocked with 5% milk in tris-buffered saline with 0.1% Tween (TBS-T, Boston Bioproduct) for 1 h. Membranes were incubated overnight with primary antibodies at 4 °C (Supplemental Table 7). Blots were washed and incubated with secondary antibodies for 2 h at room temperature. After washing, bands were visualized with ECL chemiluminescence reagents (Genesee Scientific) using the iBright Imaging System (Thermo Fisher Scientific). Band intensity was measured using the Image Studio Lite software (LI-COR Biosciences). Protein expression level was normalized to β-actin or total Tau as appropriate.

### WST-1 Assay and Neurite Length Measurement

Cell viability was measured by WST-1 reduction assay (Sigma-Aldrich). For the assay, all medium was removed and replaced with 1X WST-1 reagent dissolved in complete neurobasal medium, followed by 3 hours of incubation at 37°C. The absorbance of the culture medium was measured with a microplate reader at test and reference wavelengths of 450 nm and 630 nm, respectively. Live cell imaging was performed using the IncuCyte^TM^ Live-Cell Imaging System (Essen BioScience). Cell confluency, cell body number, neurite length, and branching points were monitored and quantified using the IncuCyte^TM^ software.

### Human iPSC-Neurons from NPC lines

Approval for work with human subjects and derived iPSCs was obtained under the Massachusetts General Hospital/MGB-approved IRB Protocol (#2010P001611/MGH). The NPC line MGH-2046-RC1 (P301L) was derived from a female individual in her 50s with FTD carrying the autosomal dominant mutation P301L (c.C1907T NCBI NM_001123066, rs63751273). The NPC line MGH-2069-RC1 (WT) was derived from a related female individual in her 40s carrying the unaffected WT Tau. Fibroblasts from the two individuals were reprogrammed into iPSCs, converted into cortical-enriched neural progenitor cells (NPCs), and differentiated into neuronal cells over 6-8 weeks by growth factor withdrawal, as previously described ^60^.

### iPSC-neurons compound treatment for western blot analysis and semi-quantitative analysis

NPCs were plated at an average density of 90,000 cells/cm^2^ of six-well plates or 96-well plates coated with poly-ornithine and laminin (POL) in DMEM/F12-B27 media and differentiated for 6 weeks. Compound treatment was performed by removing half-volume of neuronal-conditioned media from each well and adding half-volume of new media pre-mixed with the compound at 2X final concentration, followed by incubation at 37 °C. After 24h or 72h, neurons were washed in PBS, collected, and lysed. Western blot and densitometry quantifications were performed as previously described^35^.

### Tau Protein Solubility Analysis

Neuronal cell lysates and fractionation were prepared based on protein differential solubility to detergents Triton-×100 and SDS, as previously described ^61^. Briefly, cell pellets corresponding to ∼800,000 cells were lysed in 1% (v/v) Triton-×100 buffer (Fisher Scientific) in DPBS supplemented with 1% (v/v) Halt Protease/Phosphatase inhibitors (Thermo Fisher Scientific), 1:5000 Benzonase (Sigma) and 10 mM DTT (New England BioLabs). Lysates were centrifugated at 14,000 g for 10 min at 4°C. The supernatants containing Trion-soluble proteins (S fractions) were transferred to new tubes for western blot analysis. The pellets were resuspended in 5% (v/v) SDS (Sigma) in RIPA buffer supplemented with 1% (v/v) Halt Protease/Phosphatase inhibitors (Thermo Fisher Scientific), 1:5000 Benzonase (Sigma) and 10 mM DTT (New England BioLabs), and centrifugated at 20,000 g for 2 min at room temperature. These supernatants contained proteins of lower solubility/insoluble (P fractions). SDS-PAGE western blot was performed by loading 20 μg of each S-fraction and double the volume of the P-fraction onto pre-cast Tris-Acetate SDS-PAGE (Novex, Invitrogen). Western blots were performed as before. Densitometry quantification (pixel mean intensity in arbitrary units, a.u.) was done with the Histogram function of Adobe Photoshop 2022, normalized to the respective GAPDH intensity in the S-fraction, followed by normalization to Vehicle.

### Neuronal viability assays

For cardiac glycoside’s dose-dependent effects on viability, NPCs were plated (∼90,000 cells/cm^2^) and differentiated in 96-well plates for 8 weeks. After treatment with cardiac glycosides, viability was measured with the Alamar Blue HS Cell viability reagent (Life Technologies) at 1:10 dilution, after 4h incubation at 37°C and according to the manufacturer’s instructions. Readings were done in the EnVision Multilabel Plate Reader (Perkin Elmer).

For stress vulnerability assays, 1 µM or 5 µM of digoxin, oleandrin, or proscillaridin A was added to the culture media and incubated for 6h at 37 °C. Then, either 30 μM Aβ(1-42), 5 μM rotenone, 400 μM NMDA, or vehicle (DMSO) alone, was added to each well for an additional 18h of incubation. At 24h, viability was measured with the Alamar Blue HS Cell Viability reagent (Life Technologies) and the EnVision Multilabel Plate Reader (Perkin Elmer).

### Immunofluorescence of neuronal cells

NPCs were plated at a starting density of ∼90,000 cells/cm^2^ in black, clear flat bottom, POL-coated 96-well plates (Corning) in DMEM/F12-B27 media and differentiated for six weeks, followed by compound treatment. Neurons were fixed with 4% (v/v) formaldehyde-PBS (Tousimis) for 30 min, washed in PBS (Corning), incubated in blocking/permeabilization buffer [10 mg/mL BSA (Sigma), 0.05% (v/v) Tween-20 (Bio-Rad), 2% (v/v) goat serum (Life Technologies), 0.1% Triton X-100 (Bio-Rad), in PBS] for 2h, and incubated with primary antibodies overnight (Tau K9JA at 1:1000, MAP2 at 1:1000, Hoechst-33342 at 1:2500). Cells were washed with PBS and incubated with the corresponding AlexaFluor-conjugated secondary antibodies at 1:500 dilution (Life Technologies). Image acquisition was done with a Zeiss AxioVert 200 inverted fluorescence microscope.

### Data Analysis

Data management and calculations were performed using Prism 9 (GraphPad). Comparisons between two groups were done using the unpaired two-tailed student t-test. For the comparison of more than two groups, a one-way analysis of variance (ANOVA), followed by post hoc test, was performed. A P value < 0.05 was considered statistically significant, and the following notations are used in all figures: *P < 0.05, **P < 0.01, ***P < 0.001, and ****P < 0.0001. All error bars shown represent standard deviation (SD) unless otherwise stated.

